# Identification of the conserved *iol* gene cluster involved in rhizosphere competence in *Pseudomonas*

**DOI:** 10.1101/2023.05.03.538910

**Authors:** Juan J. Sánchez-Gil, Sanne W. M. Poppeliers, Jordan Vacheron, Hao Zhang, Bart Odijk, Christoph Keel, Ronnie de Jonge

**Affiliations:** Plant-Microbe Interactions, Department of Biology, Science for Life, Utrecht University, Utrecht, The Netherlands; Department of Fundamental Microbiology, University of Lausanne, Lausanne, Switzerland

## Abstract

The *Pseudomonas* genus has shown great potential as a sustainable solution to support agriculture through its plant-growth promoting and biocontrol activities. However, their efficacy as bioinoculants is limited by unpredictable colonization in natural conditions. Our study identifies the *iol* locus, a gene cluster in *Pseudomonas* involved in inositol catabolism, as a feature enriched among superior root colonizers in natural soil. Further characterization revealed that the *iol* locus increases competitiveness by inducing swimming motility and fluorescent siderophore production in response to inositol, a plant-derived compound. Public data analyses indicate that the *iol* locus is broadly conserved in the *Pseudomonas* genus and linked to diverse host-microbe interactions. Our findings suggest the *iol* locus as a potential target for developing more effective bioinoculants, given its conservation and association with diverse host-microbe interactions.

## Introduction

With a human population surpassing the 8 billion mark, modern agricultural industry faces the challenge of increasing crop yields while reducing synthetic inputs. A promising solution leverages plant-beneficial microbes as soil amendments, collectively known as bioinoculants, which have shown positive effects such as promotion of plant growth, competition against phytopathogens, and activation of host immune defenses (Bakker et al., 2020). Typically, bioinoculants need to successfully colonize the rhizosphere, which comprises the roots and the region of the soil under their direct influence, to provide such benefits. Numerous candidates for bioinoculants have been identified and studied *in vitro* or in semi-controlled conditions, and the molecular mechanisms behind their positive effects is an actively studied topic. However, the natural rhizosphere environment presents significant differences that make effective application of bioinoculants in agriculture difficult. Successful colonization of the rhizosphere depends on edaphic factors, -factors such as soil chemistry and structure-, host genotype, and, particularly, the complex web of microbial interactions in the rhizosphere microbiome that results in a strong resistance to invasion (Poppeliers et al., 2023). Consequently, many promising bioinoculants fail to exert their positive effects or show variable efficacies in the field. Nonetheless, many organisms commonly colonize and persist in the rhizosphere driven by their intrinsic capabilities, some reaching higher abundances than others. For example, Proteobacteria and Bacteroidetes are repeatedly found to be enriched within the rhizosphere of diverse plant species (Ling et al., 2022). Particularly, the *Pseudomonas* genus comprises many representatives of rhizosphere-competent organisms that display a vast diversity of competence traits and functional plasticity that grants them the ability to inhabit the rhizosphere of many plant species, like secretion systems, evasion of plant immune defenses, or production of antimicrobial compounds (Bernal et al., 2017; Liu et al., 2018; Passarelli-Araujo et al., 2022; Pieterse et al., 2021; Yu et al., 2019; Zboralski and Filion, 2020). Characterizing the underlying genetic determinants of such rhizosphere-competent microbes can help build a mechanistic understanding of the requirements of enhanced colonization and in turn, such information can be used to design future effective bioinoculants to be applied in the field.

With the aim of finding genetic features for rhizosphere-competence in natural conditions, we here compared a set of diverse *Pseudomonas* isolates from our in-lab collection for their ability to colonize the rhizosphere and root surface of *Arabidopsis thaliana* in natural soil. Genomics analysis of the best colonizers highlighted metabolism of *myo*-inositol (inositol), enabled by the *iol* locus, as an enriched explanatory feature for enhanced root colonization. To determine the contribution of the *iol* locus to root colonization we generated a *iol* deletion mutant and determined its competitiveness in *in vivo* root colonization assays in soil. We show that the *iol* mutant is less competitive and that, as expected, the *iol* locus is required for consumption of inositol. We further provide evidence suggesting that besides a nutrient source, inositol acts as a signal that induces swimming motility and production of fluorescent siderophores. Genomic analyses show that the *iol* locus is broadly conserved in the *Pseudomonas* genus, and we discuss evidence of its expression and relevance in interaction with diverse hosts and successful colonization. Together, our findings suggest a relevant role for the *iol* locus and plant-derived inositol in driving rhizosphere competence in pseudomonads.

## Results

### Competitive root colonizers are enriched for inositol phosphate metabolism

To identify genetic traits among root-associated microorganisms for competitive root colonization in natural soil conditions, we introduced, and subsequently compared the abundance of, several *Pseudomonas* isolates previously isolated from plant roots in a natural soil (**Figure S1**). To aid in isolate selection, we sequenced and compared the genomes of six root-associated pseudomonads from our collection, also including several well-known root-associated isolates (**Figure S2**). We chose a group of six isolates that varied in phylogenetic spectrum, source environments, and effects on host plants: *Pseudomonas simiae* WCS417 (WCS417), *Pseudomonas capeferrum* WCS358 (WCS358), *Pseudomonas protegens* CHA0 (CHA0), *Pseudomonas fluorescens* RS158 (RS158), *Pseudomonas* sp. WCS317 (WCS317) and *Pseudomonas* sp. WCS134 (WCS134). We introduced these isolates individually into natural soil reaching an inoculum density of 10^8^ colony forming units (CFUs)/g of soil and grew *Arabidopsis thaliana* ecotype Col-0 seedlings on this soil for 21 days. Recognizing that inference of the colonizing ability of root-associated microorganisms could be influenced by the method used to harvest the plant roots, we separated samples in two fractions, the rhizosphere which includes soil attached to the root and that is under the influence of the root, and the rhizoplane or root fraction which consists of the root without adhering soil. Following extraction of these fractions we quantified the density of each isolate in all samples by amplicon-based sequencing of the *16S rRNA* gene, using an internal control to obtain absolute abundances per sample.

Isolates displayed differential population densities across fractions (**Figure 1**, **Figure S3**). With the exception of WCS134, isolates survived similarly in bulk soil, although WCS358 and CHA0 showed slightly lower populations. Colonization in rhizosphere samples is in general lower than in soil, although the relative differences seen among isolates is very similar to the one in soil. Root samples proved to be the most effective wash for assessing differential colonization among the isolates (**Figure 1**). Specifically, WCS417 and CHA0 exhibited approximately 8.6- and 20-fold higher colonization levels, respectively, compared to the rest of isolates. We expected each isolate to show intermediate abundances in the rhizosphere fraction compared to soil and roots. However, WCS417, CHA0, and WCS134 reached higher populations in root than in rhizosphere samples, while RS158, WCS317, and WCS358 exhibit higher abundances in the rhizosphere fraction, suggesting differing lifestyles among these pseudomonads (**Figure S4**).

**Figure 1.**
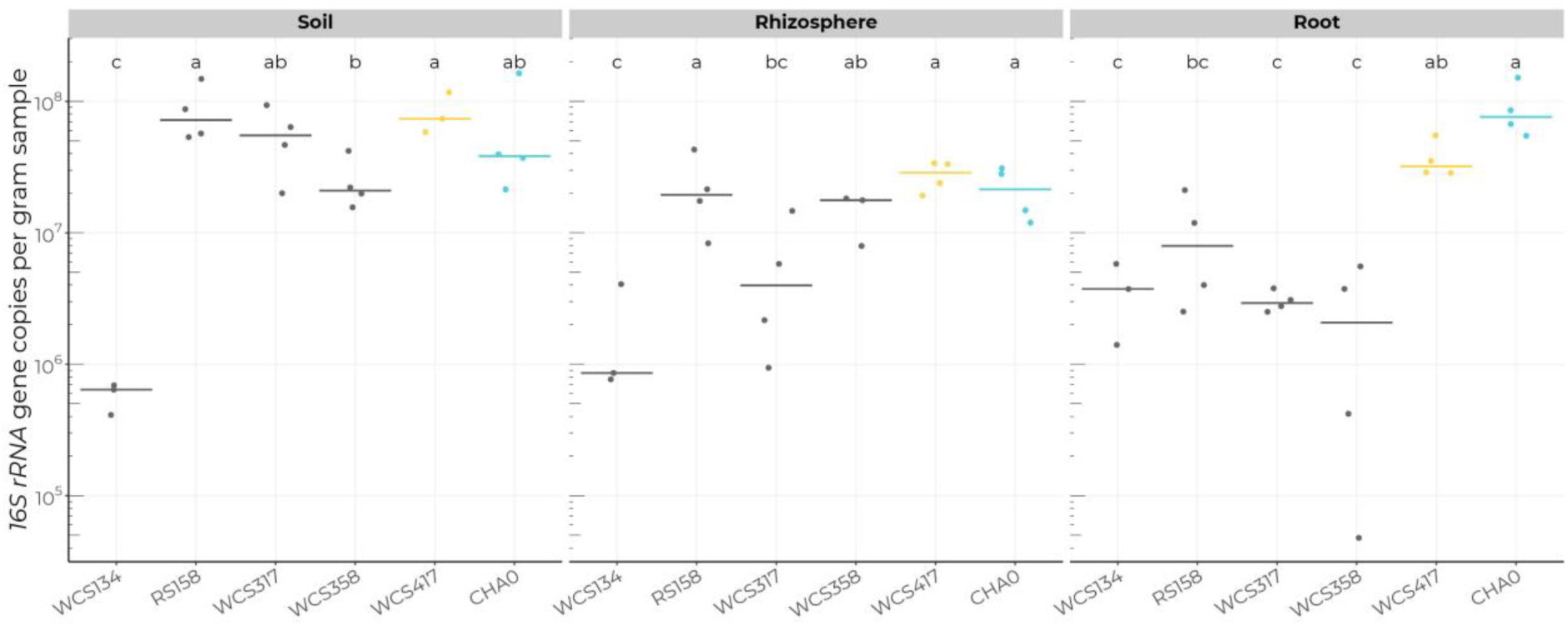
*P. protegens* CHA0 and *P. simiae* WCS417 are the best root colonizers. Absolute inoculum densities after 21 days post inoculation (dpi) measured as the number of *16S rRNA* copies per gram of sample found in soil (left), rhizosphere (middle), and root samples (right). The letters indicate groups of statistically different densities in each compartment according to a Sidak post-hoc test on negative binomial tests. *P. protegens* CHA0 and *P. simiae* WCS417 are highlighted in cyan and yellow, respectively, to differentiate them from the remaining isolates.

### Based on these results, we selected WCS417 and CHA0 as the best root colonizers and proceeded to investigate the genetic traits underlying their superior root colonization

To investigate the genetic traits that distinguish these organisms from others, we extracted the orthologous protein families shared by WCS417 and CHA0. In total, we identified 203 such families exclusively found in the best colonizers (**Figure 2a**), comprising 204 and 206 orthologues in WCS417 and CHA0, respectively. Biosynthesis of antibiotics and biofilm formation were the most common annotations in these genes, accounting for approximately 5% of the orthologues each (12 genes in WCS417 and 10 genes in CHA0 for biosynthesis of antibiotics, and 10 genes in each microbe for biofilm formation). Notably, enrichment analysis on the annotations identified metabolism of inositol phosphate, the fourth most common annotation, as the most enriched function, followed by the tenth most common annotation for biosynthesis of siderophores (**Figure 2b**). The strong enrichment for inositol phosphate metabolism is caused by the presence of a locus (hereafter *iol* locus) in WCS417 and CHA0 that is absent in the other isolates (**Figure 2c**). The locus spans 14 and 13 kbp in WCS417 and CHA0, respectively, and comprises 10 genes: a transcription factor (*iolR*), six genes responsible for inositol catabolism (*iolCEBIDG*) and three genes that encode an ABC transporter responsible for inositol intake (*iatABC*). In WCS417, an extra gene, *mocA*, encoding an oxidoreductase, is located upstream of the ABC transporters.

**Figure 2.**
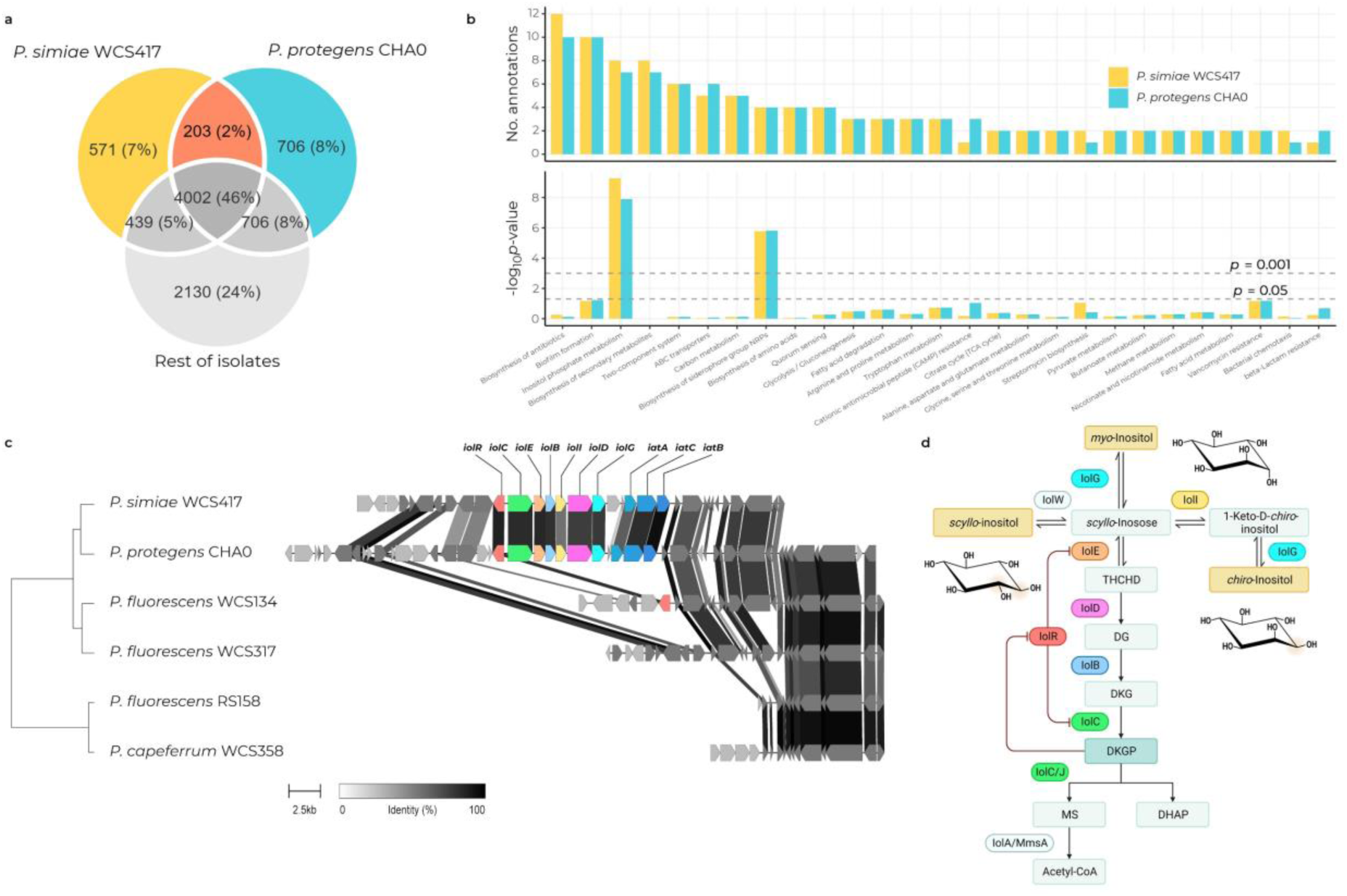
Inositol phosphate metabolism is the most enriched function in the best root colonizers. (a) Venn diagram depicting the genetic relationships between the best root colonizers and the rest of isolates. The orange area shows the set of orthologous protein families present in the genomes of WCS417 and CHA0 but absent in the other pseudomonads. (b) Bar plots showing the counts for KEGG pathways annotations (left) and their FDR-adjusted *p*-value in a Fisher’s enrichment test for the set of shared orthologues between WCS417 and CHA0, versus the other pseudomonads in this study. Only the annotations that are found more than once are shown. Note that the most enriched function is inositol phosphate metabolism. (c) Phylogenetic tree of the isolates in this study and the *iol* locus responsible for the enrichment in the annotations for inositol phosphate metabolism. (d) Predicted pathway for inositol metabolism in *Pseudomonas*. The proteins in the pathway are coloured as the genes in c. The chiral positions (C1, C2) that distinguish the *myo*, *scyllo*, and *chiro* enantiomers of inositol are shown as a shaded area on the 3D structural representations. THCHD: 3,5/4-trihydroxycyclohexa-1,2-dione; DG: 5-deoxy-glucuronate; DKG: 2-deoxy-5-keto-D-gluconate; DKGP: 2-deoxy-5-keto-D-gluconate-6-phosphate; MS: malonic semialdehyde (3-oxopropanoate); DHAP: dihydroxyacetone phosphate. Pathway in panel d was created with BioRender.com.

The IolCEBIDG enzymes potentially catalyze five consecutive reactions that transform the sugar alcohol *myo*-inositol successively into *scyllo*-inosose (IolG), trihydroxycyclohexadione (IolE), deoxyglucuronate (IolD), deoxy-keto-glucuronate (IolB), deoxy-keto-gluconate-6-phosphate (DKGP; IolC), and malonate semialdehyde and dihydroxyacetone phosphate (DHAP; IolC) (**Figure 2d**), which could theoretically enter other pathways or be directed towards anabolism as acetyl-CoA in a reaction catalyzed by MmsA/IolA, which is codified elsewhere in both pseudomonad genomes. The predicted transcriptional repressor of the locus, IolR, is inhibited by the interaction with the intermediate product DKGP, allowing the expression of the entire locus (Dong et al., 2020; Hellinckx et al., 2017; Kohler et al., 2011).

### The *iol* locus enhances competence in natural soil

To investigate the contribution of the *iol* locus to pseudomonad competence in soil, we created the multi-gene Δ*iolRCEBIDG* mutant (Δ*iol*) in CHA0 and subsequently assessed its plant root colonization. We inoculated GFP-tagged CHA0 wild type (WT) cells and mCherry-tagged Δ*iol* mutant cells individually, and a 1:1 mixture of both genotypes, in soil as before, reaching a inoculum density of 10^8^ CFU/gr of soil and then grew Arabidopsis seedlings for 21 days on this soil. In individual inoculations, colonization ability of both genotypes in soil and on the plant root was equivalent (**Figure 3a**). However, when co-inoculated, WT cells dramatically outcompeted mutant cells, achieving a 5.3-fold higher median density on the roots (**Figure 3b**). Interestingly, although the mutant achieves comparable densities when inoculated individually, its density on the root in competition was the same as in soil, suggesting that the root was primarily colonized by WT cells. As expected, the WT:Δ*iol* ratio on the root indicates a clear plant-driven advantage of WT cells (**Figure 3c**).

**Figure 3.**
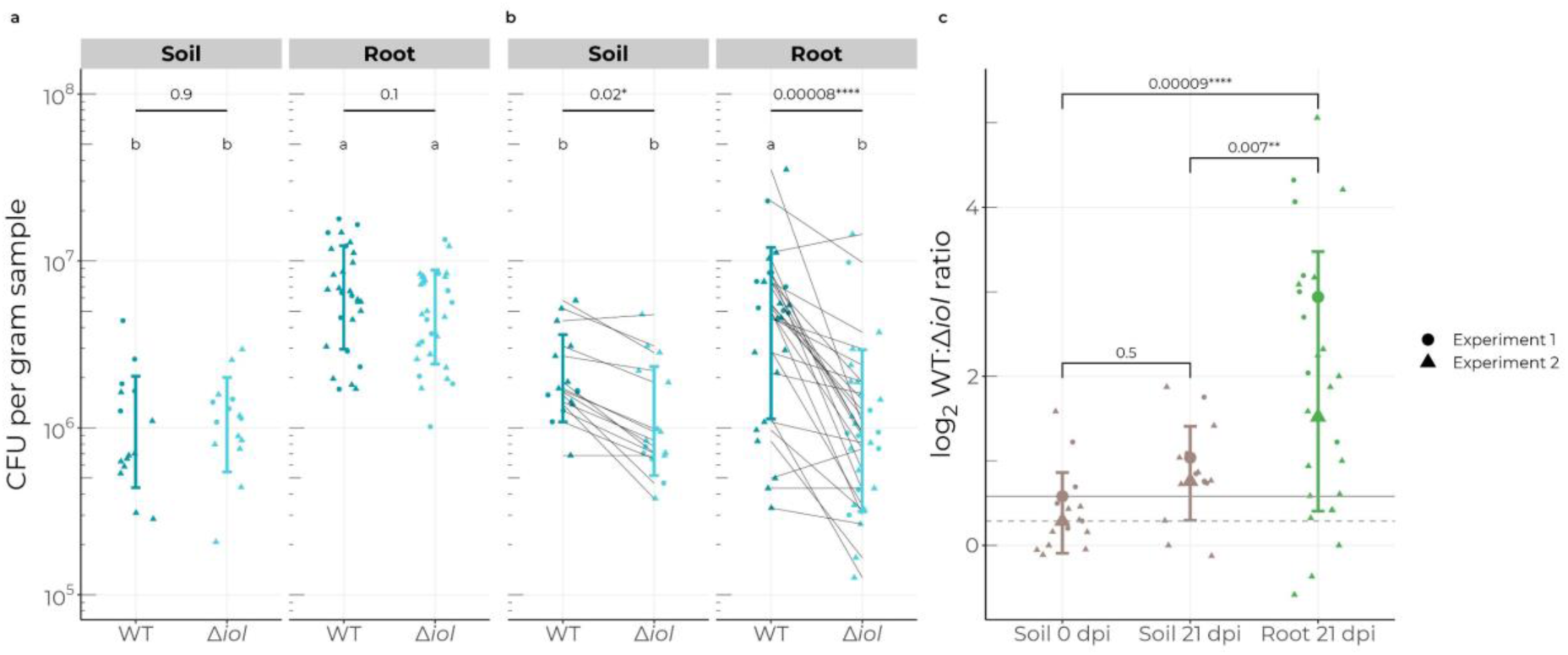
The *iol* locus enhances root colonizing competence in *P. protegens* CHA0. (a) CFU counts per gram of sample of WT and mutant cells in individual inoculations in soil (left) and root samples (right) at 21 dpi. Significance values derive from a negative binomial test for count data, adjusting for variations between experiments. (b) Paired CFU counts of each genotype in the mixed 1:1 co-inoculation at 21 dpi in soil (left) and roots (right). Significance is calculated using a paired *t*-test on all pair-wise comparisons. The letters on panels a and b indicate groupings according to a negative binomial model adjusted for experimental variation. Notice that the number of Δ*iol* cells in co-inoculation experiments are similar on roots and in soil. (c) Log_2_-transformed ratio of WT and Δ*iol* mutant cells in the mixed treatment. Large shapes indicate the median ratio per experiment, while lines show the mean initial ratio per experiment (solid line: experiment 1; dashed line: experiment 2). On the roots, WT cells achieve a significantly higher population 21 dpi, reaching an approximate ratio of 5:1.

### The *iol* locus is required for catabolism of inositol

To confirm the contribution of the *iol* locus to inositol catabolism, we performed growth curve measurements using inositol as the sole carbon source in minimal, M9, medium. Our results showed that Δ*iol* mutant cells did not exhibit any growth after 80 h, in contrast to WT cells which could utilize inositol to grow, albeit to only half the density reached with glucose (**Figure 4a-b).**

**Figure 4.**
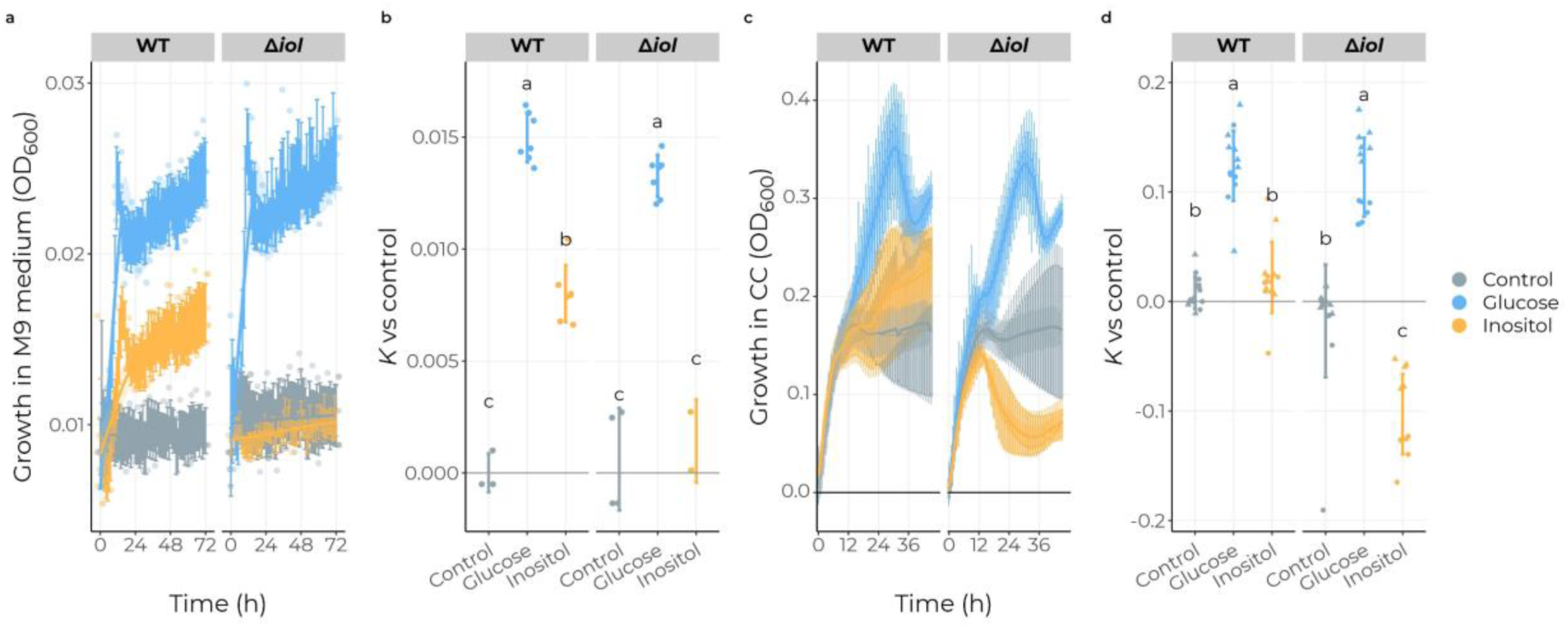
The *iol* locus is required for using inositol as a sole carbon source and affects growth of Δ*iol* cells on inositol. (a) Growth curves of WT and Δ*iol* mutant cell populations in non-supplemented minimal M9 medium (control), or in M9 medium supplemented with 100 mM glucose or inositol. Mutant cells are unable to grow with inositol, indicating that the *iol* locus is required to catabolize inositol. (b) Fitted maximum population size (carrying capacity, *K*) of the growth data in panel a. (c) Growth curves in non-supplemented CC medium (control), or in CC medium supplemented with 10 mM glucose or inositol. The data summarizes two independent growth curve assays. Despite the variability within treatments across experiments, the curves show that both genotypes stabilize in the stationary phase in control medium. Mutant cells growing in inositol-supplemented medium follow a similar pattern during the stationary phase (0-12 h), but then the optical density drops noticeably. (d) Fitted carrying capacity (*K*, maximum population size) of the growth data in panel c.

In non-minimal, CC, medium supplemented with inositol, we observed a comparable growth curve dynamic in WT cells after 36 h (**Figure 4c-d**), although the growth benefit was transient and variable (**Figure S5**). Surprisingly, we observed that inositol induced a decline in the density of Δ*iol* mutant cells after the exponential phase, at around 12 h, suggesting that inositol may have a toxic or bacteriostatic effect on these cells.

### Inositol enhances swimming motility through the *iol* locus

Inositol metabolism has been linked to motility and other relevant root colonization traits in previous studies (Fry et al., 2001; Hamilton et al., 2021; Kohler et al., 2010; Vílchez et al., 2020). Based on this, we hypothesized that the enhanced competitiveness of CHA0 in soil may also be due to an induction of motility by inositol. To test this hypothesis, we conducted typical bacterial swimming and swarming assays as we did previously for spontaneous root-competitive *P. protegens* CHA0 mutants (Li et al., 2021) in the absence and presence of inositol. These experiments revealed that inositol improved swimming motility of WT cells by 30% after 48 h while it decreased swimming ability of mutant cells by 30% (**Figure 5**). Intriguingly, we observed that while WT cell swarming motility is unaffected by the addition of inositol, mutant cell swarming is totally abolished in this condition (**Figure S6**). Glucose supplementation only promoted swimming at a 10-fold higher concentration, although the mean radius was still 10% smaller than that observed with inositol. This finding, together with the reduced growth benefit of inositol, suggests that inositol likely functions as a signal in addition to, or rather than, that of an advantageous carbon source.

**Figure 5.**
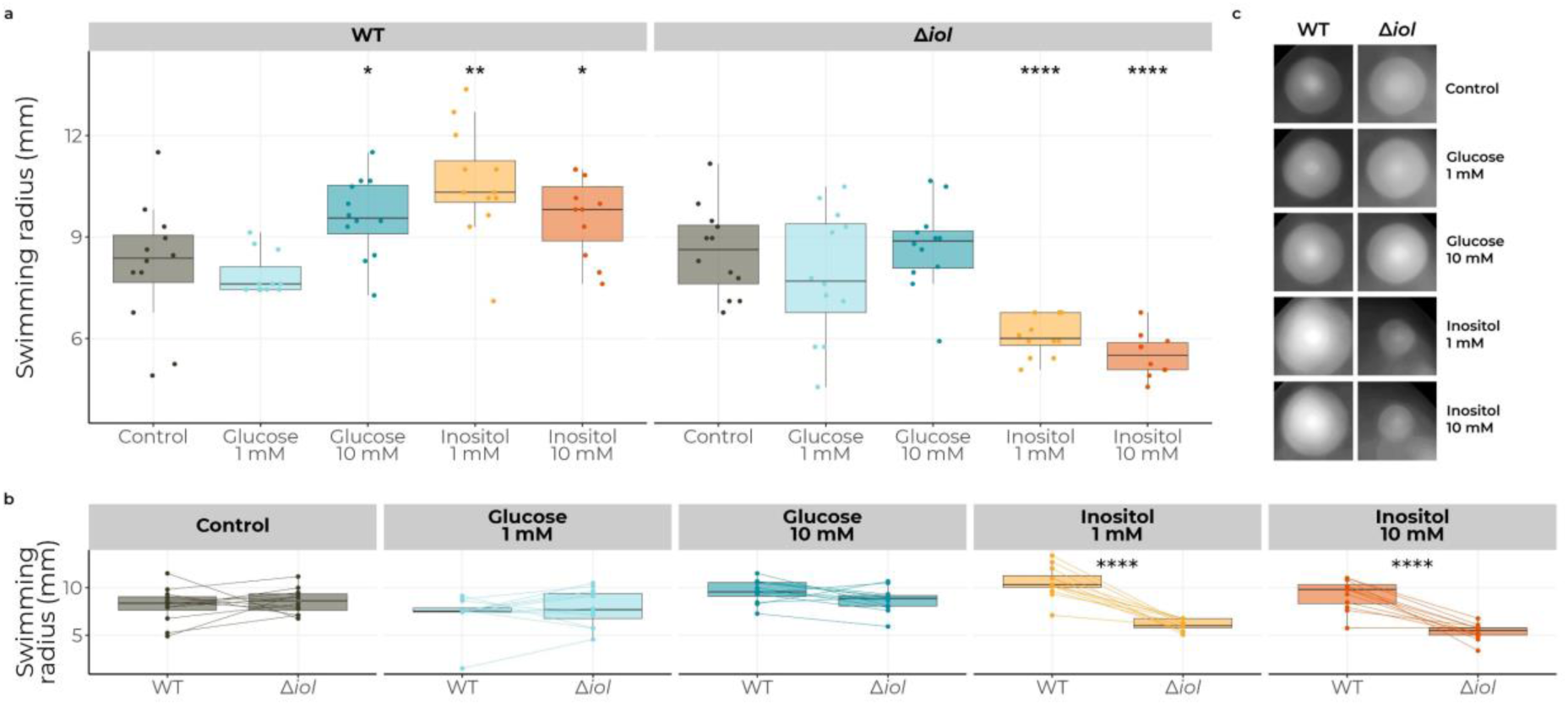
Inositol enhances swimming motility through the *iol* locus in *P. protegens* CHA0. (a) Swimming radius of WT (left) and *Δiol* mutant cells (right) in CC medium either unsupplemented or supplemented with glucose or inositol at 1 or 10 mM. Inositol induces swimming motility at 1 mM in WT cells, while motility is suppressed in the mutant. (b) Same data as in a, but shown per treatment and pairing the swimming halo for both genotypes per plate, highlighting the genotype-dependent differences in motility. (c) Representative images of WT and *Δiol* mutant cell swimming behavior in each treatment.

Besides the effect on swimming, we noticed that mutant colonies exhibited a fainter color, analogous to the decrease in optical density during planktonic growth at the onset of the stationary phase in growth curve assays. Therefore, we decided to monitor the changes in its phenotype over time in a dedicated assay. Here, we performed multiple inoculations again in swimming plates, inoculating both genotypes either individually or in combination. Mutant cells growing with inositol were visibly similar to the WT at 1 dpi, but the color intensity of the colonies faded after reaching the end of the plate, approximately at the end of the second day (**Figure S7**). Intriguingly, we observed scattered mutant colonies reappearing in plates with inositol at 6 dpi, but only in plates where the mutant grew individually (**Figure 6a**). Moreover, WT colonies at this time exhibited visibly brighter yellow colonies, indicative of an increased production of fluorescent siderophores (**Figure 6**). Altogether, these results suggest a role of the *iol* locus and inositol metabolism in quorum behavior and competence in the rhizosphere.

**Figure 6.**
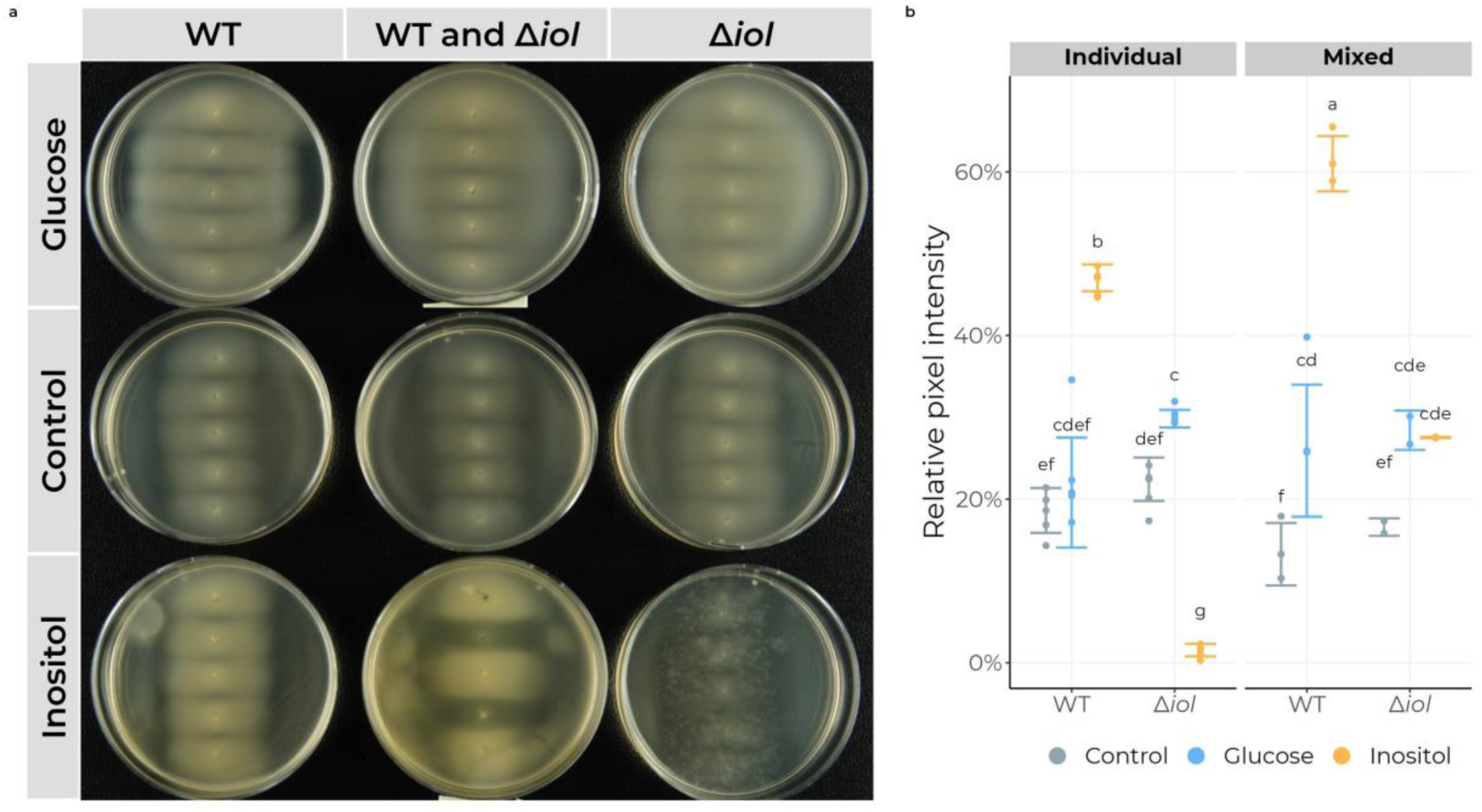
Inositol enhances the production of fluorescent siderophores and affects *Δiol* mutant physiology. Qualitative assessment of the effect of inositol and glucose on WT and *Δiol* mutant cells. (a) Representative images of colonies at 6 dpi in swimming plates supplemented with 10 mM glucose (first row), unsupplemented (second row), or supplemented with 10 mM inositol (third row). The first and third column show 5 inoculations of WT and mutant cells, respectively. The second column shows a combined inoculation of mutant cells surrounded by WT cells, used to evaluate possible interactions between both genotypes. Note how WT colonies show a brighter yellow color, characteristic of fluorescent siderophore production. Mutant colonies disintegrate at 3 dpi, and colonies re-appear at 6 dpi as seemingly independent and scattered populations. (b) Qualitative measurement of siderophores produced in the plates shown in panel a using the intensity of yellow pixels as a proxy.

### The *iol* locus is broadly conserved among pseudomonads and a determinant for diverse host-microbe interactions

Considering the potential of the *iol* locus in determining competitiveness of pseudomonads in the rhizosphere, we hypothesized that it would be a conserved trait in the *Pseudomonas* genus (**Figure 7**). Exploratory analysis of all 15,742 *Pseudomonas* genomes in the NCBI RefSeq database revealed that the *iol* locus is found in a majority of *Pseudomonas* species. Most notable exception is *Pseudomonas aeruginosa*, none of which encoded the *iol* locus (8,781 genomes, ∼56% of all genomes). From the remaining 6,962 genomes, approximately 67% contain six or more *iol* genes spaced less than 20 kbp from each other, or distributed over a limited number of genomic loci, yielding a total of 4,642 *iol*+ pseudomonads. Genomes in which the *iol* genes could not be found in a single locus typically had a low assembly quality (N50 < 50 kbp), and were considered *iol*+ if they met the requirement of the minimum number of six genes in a small (2-3) number of loci. In general, *pseudomonads* exhibit a binary *iol* locus presence-absence distribution at the species level. The frequency of *iol*+ strains is over 90% in 11 out of 23 species, and over 80% in 14 out of 23 species, not including the umbrella species *Pseudomonas fluorescens* and *Pseudomonas* spp. The remaining nine species contain at most 10% (*Pseudomonas koreensis*) *iol*+ representatives (**Figure 7b**).

**Figure 7.**
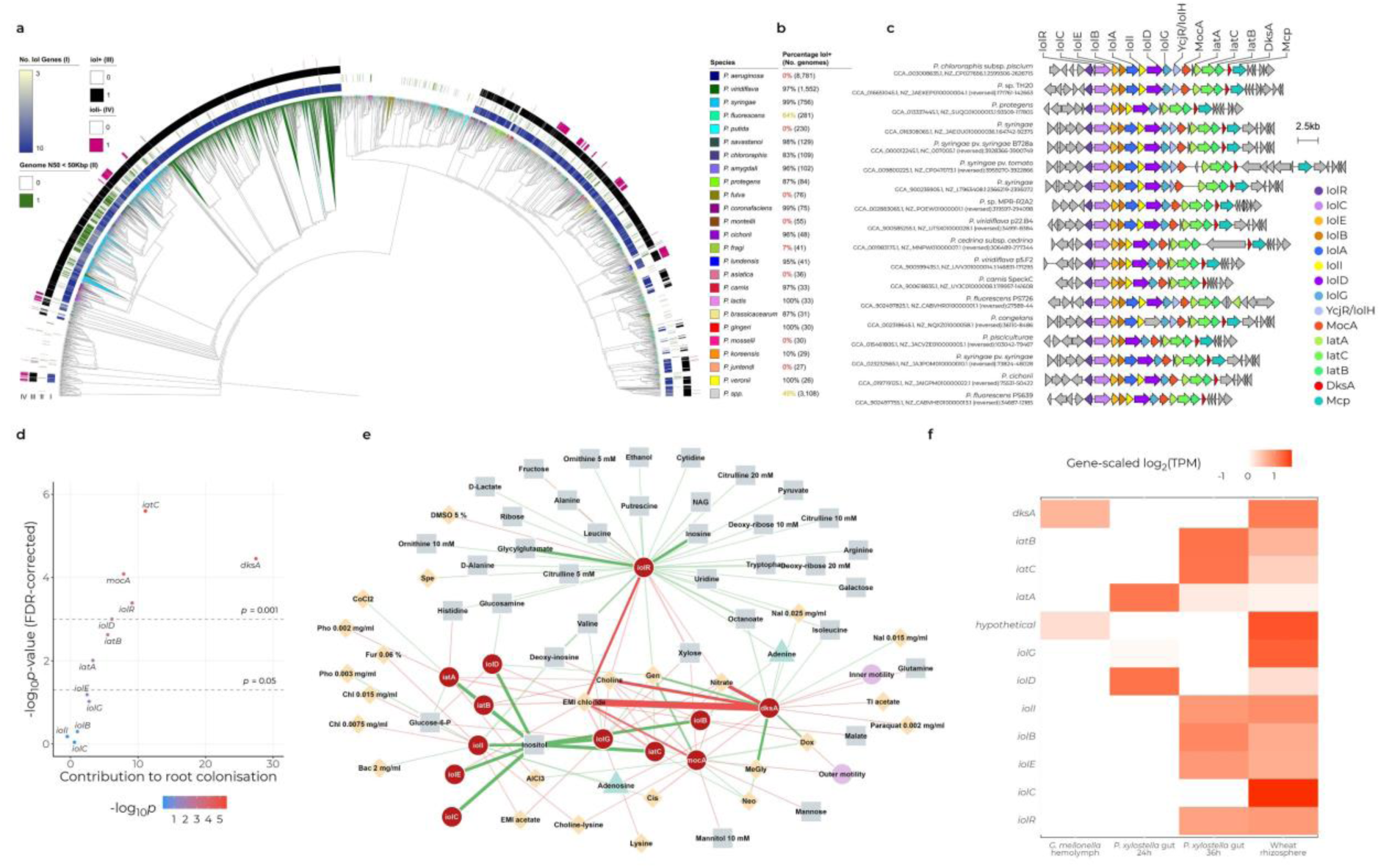
The *iol* locus is conserved in the *Pseudomonas* genus and contributes to diverse host-microbe interactions. (a) Phylogenetic tree of 15,743 *Pseudomonas* genomes available in the NCBI RefSeq database showing the distribution of the *iol* locus in the genus. 8,781 *P. aeruginosa* genomes, evolutionary distinct from the other 6,962 *Pseudomonas* genomes and all lacking the *iol* locus, were pruned from the tree. Ring I: number of *iol* genes in the highest scoring locus on the respective genome based on cblaster (Gilchrist et al., 2021). Ring II: factor indicating whether the N50 value (log2-scaled) for a given genome is below 50 Kbp as a measure for genome assembly integrity. Ring III: summary of all significantly *iol*+ scoring regions throughout the genome (minimal 6 out of 10 genes) confirming that genomes with fewer *iol* genes in the best scoring locus (ring I, light-colored), typically have a low genome N50 (ring II, green) but still encode the full locus. Ring IV: factor indicating the absence of *iolI* in the locus, showing that the loss of *iolI* follows a non-phylogenetic pattern in the genus. (b) Summary of the frequency of *iol*+ genomes in each *Pseudomonas* species, according to the annotated species names. (c) Locus alignment from a representative set of genomes in the tree, showing locus conservation in the genus. Some loci harbor an extra *iol* gene, *iolA*/*mmsA*, downstream of *iolB*, and two other genes, *mocA* and *ycjR*, encoding the sugar phosphate isomerase/epimerase *iolH*, upstream of the genes encoding the ABC transporters. (d) Scatter plot showing the relative contribution of the genes in the *iol* locus in *P. simiae* WCS417 to Arabidopsis root colonization *in vitro* based on publicly available mutant fitness data (Cole et al., 2017). The x-axis represents the contribution of each gene to colonization, using the *t*-like effect-size statistic versus initial mutant populations, as previously reported (Cole et al., 2017). The y-axis represents the FDR-corrected *p*-value for the contribution of each gene, as inverse logarithm of the raw value for ease of interpretation. (e) Bipartite network depicting the contribution of each gene in the iol locus in *P. simiae* WCS417 to survival in different environments included in the Fitness Browser (Price et al., 2018). Genes are shown as red nodes, and links connect these with conditions in which they show a differential fitness across genome-wide fitness assays. Green and red links indicate a positive or negative contribution to survival, respectively. Link width and opacity indicate link weight (absolute contribution). Conditions are colored by their nature: carbon sources (grey), stressors (yellow), or nitrogen sources (light teal). Notice how *iolR* is not directly related to survival in inositol but in a myriad of other carbon sources and specific stressors. Notably, *iol* genes show many links to antimicrobial compounds. EMI, ethylmethylimidazolium; Gen, gentamicin; Neo, neomycin; Pho, phosphomycin; Dox, doxycycline; Bac, bacitracin; Chl, chloramphenicol; Cis, cisplatin; Fur, furfuraldehyde; Nal, nalidixate; Spe, spectinomycin; DMSO, dimethylsulfoxide; CoCl2, cobalt (II) chloride; AlCl3, aluminium (III) chloride; Tl acetate, thallium acetate; MeGly, methylglyoxal. (f) Heatmap of gene expression data from (Vesga et al., 2020), showing expression of the *P. protegens* CHA0 *iol* locus during insect and plant colonization/infection. Other relevant colonization genes are also included. Expression data is shown as gene-scaled log_2_-scaled transcripts per million).

Among *iol*+ genomes, the core *iolRCEBDG* genes are found in a very high frequency, between 97% and 100% of genomes, indicative of co-evolution as a locus and suggesting that all genes are required to perform its biological function in the proposed catabolic pathway. Interestingly, around 10% of *iol*+ genomes lack *iolI* (454 pseudomonads), and the distribution of these 454 over the phylogenetic tree suggests that *iolI* loss occurred multiple times throughout the evolution of the genus. *IolI* encodes an inosose isomerase which is responsible for the conversion of 1-keto-D-*chiro*-inositol into *scyllo*-inosose. Absence of *iolI* could therefore indicate the lack of *chiro*-inositol-derived compounds in the environment of these specific organisms. Other common structures of the locus contain a copy for a methylmalonate semialdehyde dehydrogenase (*iolA*/*mmsA*) between *iolB* and *iolD*, and/or a sugar phosphate isomerase/epimerase between *iolG* and the *iatACB* genes, potentially *iolH* (**Figure 7c**). Neither of these genes was found in WCS417 and CHA0. Lastly, a gene encoding a transcription factor, *dksA*, and a putative chemotaxis protein, *mcp*, are located downstream of the *iatACB* genes, suggesting a direct connection between the *iol* locus and motility.

In order to search for more evidence of the relationship of root colonization and the *iol* locus, we explored publicly available transposon-insertion sequencing (Tn-Seq) data in WCS417 (Cole et al., 2017). In this study, a collection of random insertion WCS417 mutants was let to colonize Arabidopsis roots in plates for seven days, after which the abundance of each mutant was used to assign a fitness (or performance) value for root colonization for each gene. According to our analysis, root colonization appears supported by *iolR*, the *iatABC* transporters, *iolD*, and the *traR/dksA* transcription activator. However, the remaining genes in the locus (*iolCEBIG*) did not show a significant contribution to colonization, suggesting that consumption of inositol is not *per se* the most determinant feature for successful colonization *in vitro* (**Figure 7d**). To further investigate the relationship between inositol and colonization in the context of the *iol* locus, we made use of publicly available data from additional genome-wide, transposon mutant-based fitness experiments in WCS417, in which mutants were exposed to many different conditions, including stresses, carbon and nitrogen sources, and motility assays (Price et al., 2018). We used this data to construct a bipartite network connecting the genes in the *iol* locus to the conditions where they showed a significant fitness contribution (**Figure 7e**). Unexpectedly, while most of the genes seemed related to metabolism of inositol, neither *iolR* nor *dksA* appear to contribute directly to fitness in inositol as a sole carbon source. Instead, our analysis revealed that *iolR* is primarily related to consumption of other carbohydrates, while *dksA* plays a role in survival when cultured with antimicrobial compounds. Both transcription factors contribute to susceptibility to stressors like nitrate, choline, and imidazolium chloride, and promote survival in gentamicin. Interestingly, we observed a negative relationship of *dksA* with swimming motility, suggesting that the induction of motility by inositol in WT cells might be due to an inositol-induced inhibition of DksA activity. We did not find any obvious connection to the production of fluorescent siderophores.

Based on the manifest connection between root colonization and the locus, we set to explore its expression during host colonization. For this, we examined publicly available data from a study performed by Vesga and co-authors (2020). Here, the transcriptional activity of *P. protegens* CHA0 was assayed during the colonization of the hemolymph of an insect host (*Galleria mellonella*), of the insect gut (*Plutella xylostella*), and of wheat roots (*Triticum aestivum*) (Vesga et al., 2020). We found that the *iol* locus is expressed during colonization on wheat roots and in the insect gut, but not in the insect hemolymph. Considering that hemolymph data was collected following direct injection, these data suggest that the *iol* locus is related to host perception and/or colonization (**Figure 7f**).

## Discussion

In our study, we compared the colonization ability of a set of pseudomonads in non-sterile natural soil by amplicon-based sequencing of the *16S rRNA* gene. The use of spike-in internal controls for absolute quantification from complex environments has been applied in recent studies (Kim et al., 2021; Tkacz et al., 2018), and permits high-throughput quantification that does not depend on isolate-specific resistance markers and/or dilution series of the samples. Using this approach, we found that the compared colonization ability in our set of isolates varied substantially between rhizosphere and root samples (**Figure 1**), which suggests that these pseudomonads exhibit different lifestyles and environmental preferences: microbes enriched on the root will be firmly attached to the host and embedded in a biofilm on the surface -i.e., WCS417 and CHA0-, while organisms preferring the rhizosphere might benefit from diffusing nutrients and potentially avoid stress from an active immune system. In any case, our findings indicate that careful consideration of the method selection should be taken in microbial colonization studies, as different treatments might select for microbes with different lifestyles and lead to disparate conclusions about their compared ecological competence.

The two best root colonizers, WCS417 and CHA0, are well-known pseudomonads that have been widely studied for their plant-beneficial effects (Pieterse et al., 2021; Vesga et al., 2020). The adaptation to a plant-associated lifestyle requires an optimized consumption of plant-derived resources, among which inositol and its derivatives play an important role. In plants, inositol polyphosphates are main constituents of the plasma membrane and have key roles in signaling, immunity, and phosphorus storage (Castrillo et al., 2017; Dindas et al., 2022; Gomes et al., 2005; Tan et al., 2007), and inositol exudation through root tissues has been reported in many species (Chaparro et al., 2014; Pini et al., 2017). The ability to metabolize inositol emerged as a potentially distinctive trait in plant-associated pseudomonads in a genomic meta-analysis (Levy et al., 2018), supporting the idea that the presence of the *iol* locus in WCS417 and CHA0 arises indeed from optimized adaptation to plant hosts. In general, expression of the *iol* genes *in planta* or in response to root exudates has also been observed, although often indirectly or outside of the experimental focus of the study. Expression of the *iol* genes in isolates from the genera *Cupriavidus, Burkholderia*, and *Rhizobium* was upregulated in response to root exudates of the legume *Mimosa pudica* (Klonowska et al., 2018). In pseudomonads, expression of the locus and its relation to competence has also been observed in isolates with divergent lifestyles. The pathogen *Pseudomonas syringae* pv. *tomato* DC3000 expresses the *iol* genes when grown in tomato apoplast extracts (Rico and Preston, 2008), and during growth in the Arabidopsis apoplast (Nobori et al., 2018). Likewise, *Pseudomonas syringae* pv. *syringae* B728a requires a functional *iol* locus for full pathogenicity in bean leaves (*Phaseolus vulgaris*), especially during apoplastic colonization (Helmann et al., 2019; Yu et al., 2013). Notably, we found that the vast majority of *Pseudomonas syringae* genomes are *iol*+, highlighting its relevance for this plant pathogenic bacterium (**Figure 7b**). These findings are further corroborated by a recent, comprehensive analysis of the distribution of the *iol* locus by Weber and Fuchs (2022) revealing that hundreds of bacteria from diverse families and occupying disparate ecological niches encode the capacity to catabolize inositol via the *iol* locus. These include animal pathogens, plant pathogens, soil commensals and rhizosphere-associated bacteria like the here discussed pseudomonads (Weber and Fuchs, 2022).

However, the precise physiological function of the *iol* locus seems to be variable and isolate-specific. In a study comparing expression patterns of a selection of eight pseudomonads in response to root exudates from *Brachypodium distachyon*, six out of eight strains showed up-regulation of *iol* genes. *Pseudomonas protegens* Pf-5, the closest relative of CHA0, increased the expression of the entire locus and yet did not exhibit any growth when grown in inositol as a sole carbon source, suggesting again that utilization of inositol for growth is not the only function of the locus. In fact, up-regulation of the *iol* locus in the other pseudomonads in this study, either fully or partially, did not seem to be a requirement for growth in inositol at all.

In our root colonization assays, we show that Δ*iol* mutant CHA0 cells can colonize equally to WT cells when inoculated individually, while they fail in reaching a higher population density than in soil when they compete with WT cells. We also observed growth defects in mutant cells when grown in inositol. Interestingly, a similar phenomenon has been described in the Alphaproteobacteria *Sinorhizobium meliloti*, *Sinorhizobium fredii*, and *Rhizobium leguminosarum*, where the *iol* genes play an important role in defining competence in root nodules (Fry et al., 2001). In *Sinorhizobium*, *iolG* mutants form abnormal bacteroids, are impaired in nitrogen fixation, and are unable to utilize rhizopines, *scyllo*-inositol derivatives produced in nodules, which ultimately results in a loss of competitiveness against WT cells (Galbraith et al., 1998; Jiang et al., 2001; Kohler et al., 2010; Murphy et al., 1995). Yet, these growth defects do not seem to pose a disadvantage in our individual inoculations, as mutant density on the root is not different from WT populations. The stimulation of swimming motility in WT cells by inositol, however, might promote early arrival during initial stages of colonization and therefore enable a quicker occupation of the root surface. In the plant pathogenic bacterium *Ralstonia solanacearum*, expression of the *iol* genes is restricted to the initial onset of infection by a quorum-sensing mechanism. Knock-out mutants for *iolG* in *R. solanacearum* are impaired in their capacity to colonize tomato roots during the initial stages of infection (Hamilton et al., 2021), and, remarkably, swimming motility during this phase has also been linked to successful pathogenicity in an independent study (Corral et al., 2020). A direct link between inositol and bacterial motility was previously observed in *Priestia megaterium* (formerly *Bacillus megaterium*). In this organism, inositol exuded by the roots of Arabidopsis and tomato plants (*Solanum lycopersicum*) induced a strong chemotactic response and enhanced biofilm formation (Vílchez et al., 2020). Noteworthy, quorum-dependent responses, like the one described in *Ralstonia* and *Priestia*, commonly take place in the transition to the stationary phase (Goo et al., 2012). Furthermore, rhizopines, which act as quorum-sensing signals in *Sinorhizobium* species (Zuniga-Soto et al., 2019), affect motility, growth, and biofilm formation, and also act during the stationary phase and not before (Krysciak et al., 2014). The connection between inositol metabolism and quorum behavior could explain the decrease in optical density in the Δ*iol* CHA0 at the onset of the stationary phase that we observe, and, additionally, the enhanced production of siderophores that we observed after six days on plates.

Our findings imply an important role for the *iol* locus during *Pseudomonas* root colonization beyond utilization of inositol as a carbon source. Our analyses of publicly available genome-wide fitness data show that root colonization *in vitro* depends primarily on the *iolR*, *iatABC*, and *dksA* genes, while being seemingly independent of inositol consumption. Additionally, IolR does not affect inositol consumption, contrasting with previous results (Dong et al., 2020; Hellinckx et al., 2017; Kohler et al., 2011), but is required for root colonization and exhibits many links to metabolism of diverse carbon sources. Moreover, swimming motility appears to be inhibited by DksA, suggesting that the induction of swimming by inositol in our experiments might be due to an inositol-induced repression of DksA. Together with our findings on the overproduction of fluorescent siderophores, all this information supports a more complex mechanism in response to the presence of host-derived inositol in the environment. Indeed, IolR regulates behavioral responses towards inositol in other organisms, like cell autoaggregation and biofilm formation in *Aeromonas hydrophila* (Dong et al., 2020), the expression of type III secretion system in *Salmonella typhimurium* (Cordero-Alba et al., 2012), or xylose uptake in *Corynebacterium glutamicum* (Klaffl et al., 2013).

In general, the *iol* locus has been associated to a myriad of different phenotypes in other bacteria, and its physiological and ecological functions also seem to go beyond mere obtention of energy from inositol. Instead, the locus appears to be embedded in a complex mechanism of sensing and response to host-related environmental cues. For example, in *Caulobacter crescentus* the iron-deficiency response regulator Fur represses expression of the *iol* locus in an iron-independent manner (da Silva Neto et al., 2013), and in *Salmonella typhimurium*, the locus is linked to antibiotic resistance (Lynch and Kierstead, 1985). In salmonellae, the small RNA *RssR* controls expression of the *iol* locus and inositol metabolism (Kröger et al., 2018), leading to complex expression dynamics controlled at the bacterial population level (Kröger et al., 2011).

The specificity of the phenotype associated to the *iol* locus in pseudomonads may be reflected in the diversity of locus decorations. Our analysis of the *iol* locus among available *Pseudomonas* genomes shows that besides the core locus (*iolRCEBDG*), specific conformations with additional genes can be found. Particularly, we observed that *iolI* has been lost in ∼10% of *iol*+ pseudomonads. The inosose isomerase IolI enables uptake of *chiro*-inositol through the conversion of 1-keto-D-*chiro*-inositol into *scyllo*-inosose, which can be readily incorporated into the main pathway. Although derivatives of inositol can be found in soil in diverging amounts, the *chiro* conformation is rare in soil. However, it is produced by plants of many species in differing amounts and forms (Gomes et al., 2005; Siracusa et al., 2022; Turner et al., 2002; Xia and Wang, 2006), which further supports the link between the presence of *iolI* with a plant-associated lifestyle. Together with the lack of well-defined taxon-specific loss of *iolI*, this suggests that the different structures of the *iol* locus that we described here might correspond to host-specific locus conformations according to the composition of host exudates. Divergent *iol* loci in *Pseudomonas* can include several other genes, like *iolA*, *mocA*, and *dksA*. The methylmalonate semialdehyde dehydrogenase IolA/MmsA produces acetyl-coenzyme A (CoA) from the last digestion product in the predicted pathway, malonate semialdehyde, or malonyl-CoA from methylmalonate semialdehyde. This gene is frequently found elsewhere in the genome, suggesting that this conversion can be regulated in both inositol-dependent and independent ways. The *mocA* gene was originally annotated as a Gfo/Idh/MocA-like oxidoreductase, which is a large protein family with many disparate functions described, among them the metabolism of rhizopines in *Sinorhizobium* species (Taberman et al., 2016). In parallel, annotations of this gene in some pseudomonads suggest that this gene might encode a homolog of IolW, a *scyllo*-inositol dehydrogenase that catalyzes conversion of this inositol enantiomer into *scyllo*-inosose and allows its incorporation in the main pathway. Both annotations, MocA and IolW, point towards the use of *scyllo*-inositol, or rhizopine derivatives, and possible associations with nodulating bacteria. Further experiments are required to clarify whether pseudomonads can in fact use rhizopines or other *scyllo*-inositol-derived exuded compounds in the rhizosphere. Lastly, DksA, encoded in the 3’-end of the locus, appears involved in survival in stress induced by methylglyoxal in our analysis of the publicly available genome-wide fitness data. This is one of the few reported phenotypes for IolS in *Bacillus* species, an aldo-keto reductase with otherwise unknown function (Chandrangsu et al., 2014; Ehrensberger and Wilson, 2004). Like DksA in pseudomonads, IolS does not seem to be required for inositol metabolism, yet it is conserved in the *iol* locus in bacilli and up-regulated in response to maize exudates (Fan et al., 2012).

In conclusion, we show that the *iol* locus determines rhizosphere competence in pseudomonads. Detailed characterization further revealed that its function entails more complex mechanisms than mere consumption of the sugar alcohol *myo*-inositol, and that these mechanisms are taxon- and possibly host-specific. Future research should dig deeper into the mechanistic processes by which competence is optimized by the *iol* locus in the *Pseudomonas* genus. The findings in our study provide evidence of the fundamental basis of rhizosphere colonization and signify an important step towards development of robust bioinoculants in sustainable agriculture.

## Acknowledgements

The authors want to thank Bas E. Dutilh, Corné M.J. Pieterse and members of the Plant-Microbe Interactions lab for helpful discussions. We acknowledge the Utrecht Sequencing Facility (USEQ) for providing sequencing service and data. USEQ is subsidized by the University Medical Center Utrecht and The Netherlands X-omics Initiative (NWO project 184.034.019). This study was supported by the NWO Green II Grant no. ALWGR.2017.002 (S.W.M.P., J.J.S.G., R.d.J.), the Novo Nordisk Foundation Grant no. NNF19SA0059362 (R.d.J.), and the China Scholarship Council scholarship no. 201406300090 (H.Z.).

## Author contributions

**Juan J. Sánchez-Gil:** Conceptualization, Methodology, Software, Validation, Formal analysis, Investigation, Data curation, Visualization, Writing – Original Draft. **Sanne W. M. Poppeliers, Bart Odijk:** Investigation. **Jordan Vacheron and Christoph Keel:** Methodology, Resources, Writing – Review and Editing. **Hao Zhang:** Methodology, Resources. **Ronnie de Jonge:** Supervision, Project administration, Funding acquisition, Validation, Formal analysis, Writing – Review and Editing.

## Materials and methods

### Arabidopsis culture

Seeds of Arabidopsis accession Col-0 were surface-sterilized with chlorine gas in a glass chamber and subsequently aerated in a flow cabinet to remove the residual chlorine for 30 min. Sterile seeds were sown in agar plates containing 1x Murashige and Skoog (MS) medium supplemented with 0.5% sucrose. Plates were sealed twice with parafilm, let stratify for 48 h at 4°C, and positioned vertically in a growth chamber under short-day conditions (22 °C; 10 h light/14 h dark; light intensity 100 μmol/m^2^/s).

### Soil treatments, transplantations, and microbiome sampling

Per isolate, single colonies were spread onto five King’s B (KB) medium agar plates and incubated overnight at 28 °C. Per plate, 5 ml of MgSO_4_ 10 mM was used to scrape the microbial cell layer off the surface, and the resulting suspension was collected into 50-ml tubes for every isolate. The cell suspension was washed thrice by centrifugation at 3000 x g for 10 min, resuspending in 40-50 ml MgSO_4_ 10 mM every step. The final cell pellet was resuspended in 20 ml of MgSO_4_ 10 mM, and the bacterial density was measured by determining the optical density (OD) at 600 nm (equivalence: 1 OD_600_ unit, 8·10^8^ CFU/ml).

Each microbial suspension was added to dried Reijerscamp soil to reach an inoculum density of 10^8^ CFU/g of soil, with a final volume of 100 ml/kg of soil. For the mock treatment, the soil was moistened with MgSO_4_ 10 mM at 100 ml/kg of soil.

Per inoculation treatment, 12 two-weeks-old Arabidopsis seedlings were transferred to individual pots (diameter = 5.5 cm and height = 5 cm; MXC5,5 Pöppelmann) containing 100 g of inoculated Reijerscamp soil each. All pots were placed on trays on individual Petri dishes to prevent cross-contaminations and kept in the growth chamber under the same conditions as mentioned above. The trays were kept closed with a lid for the first two days post-transplantation. For the remaining time of the cultivation period, the lids were removed, and the plants were watered every two days with 5-10 ml of tap water. Roots were eventually harvested at 21 dpi (5 weeks-old) and shaken to remove most attached soil particles. For rhizosphere samples, roots were placed in 50-ml tubes with 30 ml sterile 10 mM MgSO_4_ and the roots were washed by inverting the tube 10 times. For root samples, roots were washed twice in 50-ml tubes by vortexing 15 s in 30 ml sterile phosphate-Silwet buffer (PBS-S; 137 mM NaCl, 10 mM Na_2_HPO_4_, 1.8 mM KH_2_PO_4_, 2.7 KCl, 0.02% Silwet). Roots cleaned with both methods were retrieved with sterile tweezers, tapped dry on clean paper, and transferred to 2-ml tubes containing 1 ml MgSO_4_ 10 mM with two glass beads. For soil samples, we collected approximately 0.25 g of soil from unplanted pots. Sample weight was obtained by weighing the tubes before and after collecting the root sample. The tubes were shaken twice at 20 Hz in a TissueLyser, snap frozen in liquid nitrogen, and stored at -80°C until DNA extractions.

### DNA extractions, *16S rRNA* amplicon library preparation, and data normalization

Nucleic acid extractions were performed on a KingFisher Flex (Thermo Fisher Scientific) using a protocol based on the MagAttract PowerSoil DNA EP Kit and the DNeasy Powersoil Kit. The resulting DNA was quantified with Qubit fluorometric quantitation (Invitrogen) with the BR dsDNA assay kit (Thermo Fisher Scientific).

In order to obtain absolute counts of each isolate, we used DNA from *Salinibacter ruber* (*Salinibacter*), an extreme halophile found in saltern crystallizer ponds that is not found in our samples and that encodes a single *16S rRNA* gene copy in its genome. Based on the genome size of 3.6 Mbp (GenBank accession ID GCA_000013045.1), we calculated an expected copy number of 2.46·10^5^ chromosomes per nanogram of genomic DNA, with an equivalent number of copies of the *16S rRNA* gene. Cell lysis buffer was spiked with 4 ng of *S. ruber* DNA per sample, aiming at a final concentration of 0.1-1% of the total amount of DNA. Library preparation was conducted according to Illumina’s 16S Sequencing Library Preparation protocol, using phasing primers for the V3-V4 region (**Table S1**). The final libraries were quantified using Qubit fluorometric quantitation with the BR dsDNA assay kit, pooled at a sample concentration of 4 nM, and sequenced on Illumina MiSeq to obtain 2×300 bp paired-end reads.

The resulting reads were processed using cutadapt (Martin, 2011) and the DADA2 pipeline (Callahan et al., 2016). To increase sensitivity of the DADA2 pipeline for the introduced isolates and their respective *16S rRNA* sequences, we used the known V3-V4 region of the isolates, plant mitochondrion, plant chloroplast, and *Salinibacter* as priors in the amplicon sequence variant (ASV) inference step. For every inoculation treatment, ASVs were inferred from all treated samples and mock samples by prioring with the sequence of the respective isolate, plant, and *Salinibacter*. Sequences used as priors are listed in **Table S2**.

After removal of chimeric sequences, the counts of every isolate were normalized with respect to the counts of *Salinibacter* by taking the ratio of isolate counts to the observed *Salinibacter* counts, and multiplying this ratio by the total number of added *Salinibacter* copies in the sample preparation (4 ng/sample · 2.46·10^5^ 16S gene copies/ng, 9.84·10^5^ copies/sample). Final microbial densities are calculated by normalizing estimated *16S rRNA* copy numbers with plant weight and subtracting counts from uninoculated samples when applicable.

### Genomic analysis

To identify genes that could potentially explain the enhanced colonization ability in *P. simiae* WCS417 and *P. protegens* CHA0, we subjected the genomes of all isolates to a comparative genetic analysis based on presence/absence of orthologs. First, proteomes were inferred from the genome sequences using Prokka (Seemann, 2014). The proteomes were then used as input for OrthoFinder (Emms and Kelly, 2019) in order to infer phylogenetic relationships among all genes and group them into orthologous protein families or orthogroups. The result is a table of all orthogroups inferred and a list of its orthologues found in each individual genome. In parallel, all proteomes were annotated using the eggNOG-mapper (Huerta-Cepas et al., 2019) and the KEGG database (Kanehisa and Goto, 2000). Enriched functions in the set of orthologues shared by only WCS417 and CHA0 were identified by a Fisher’s exact test by comparing the functions of these orthologues with those in the complete proteomes.

### Mutant construction

Deletion of the entire *iol* gene cluster (from PPRCHA0_2627 to PPRCHA0_2633, i.e., from *iolR* to *iolG*) was generated using the suicide vector pEMG and the I-SceI system adapted to *P. protegens* (Heiman et al., 2022; Vacheron et al., 2021). Primers and plasmids used are listed in **Table S3**.

### Root colonization and competition assays

Experiments with *gfp*-tagged CHA0 WT and *mCherry*-tagged mutant were performed as described previously for the isolates comparisons, with minor differences. Individual inoculations were performed at the same microbial density as described, 10^8^ CFU/g soil, while mixed inoculations contained 0.5·10^8^ CFU/g of each isolate, in order to obtain a comparable total microbial density as in individual inoculations. At 21 dpi, only root and not rhizosphere samples were collected. Instead of freezing, 10-fold dilutions of the shaken supernatant were plated on KB agar plates supplemented with 10 μg/ml gentamicin and incubated overnight at 28°C. Root microbial density was calculated by normalizing CFU counts by root fresh weight.

### Growth assays

For planktonic growth, overnight cultures of CHA0 WT and the Δ*iol* mutant were diluted 1/10 in fresh, liquid KB medium and grown 4 h in order to synchronize both genotypes in the exponential phase. The cell suspension was washed thrice by centrifugation at 3,000 x g, decanting of the supernatant, and resuspension in the respective medium. Cells were inoculated individually in flat-bottom 96-well plates at a final OD_600_ equal to 0.01 in either M9 (assessment of sole carbon sources) or 0.1X Cook’s Cytophaga (CC). For assessment of inositol as a carbon source, the medium was left unsupplemented or supplemented with 100 mM of glucose or inositol. For assaying effects of inositol on growth, medium was left unsupplemented or supplemented with 10 mM glucose or inositol.

Growth curves were obtained by measuring OD_600_ every 30 min in a plate reader for 72 h (inositol usage as carbon source) or 48 h (inositol effects on growth). Growth curve analyses were done with the *growthcurver* package in R (Sprouffske and Wagner, 2016).

### Motility assays and siderophore plates

Swimming and swarming motility were assayed in 0.1X Cook’s Cytophaga-agar (CCA) minimal medium at 0.3% or 0.5%, respectively. For all assays, the medium was supplemented with inositol or glucose as a control carbon source at 1 or 10 mM or left unsupplemented. Cells were prepared as for growth experiments. For the swimming assay, sterile pipette tips were dipped in the cell suspension and the agar was gently pierced until reaching the middle of the agar, and without reaching the bottom. Pictures were taken after 48 h of growth at 21 ⁰C and analyzed with ImageJ (Schindelin et al., 2015). The swimming halo radii and diameters were used to assess swimming motility. For swarming, 1 μl of bacterial suspension was carefully inoculated on the agar surface and avoiding damage of the agar. Pictures were taken at the same time points as for swimming, and swarming ability was measured as the surface area covered by bacteria. In both assays, we inoculated CHA0 WT, CHA0 Δ*iol*, and two previously reported spontaneous CHA0 mutant that exhibit promoted swimming motility as positive control (*sadB*), and impaired swimming as negative control (*fleQ*) (Li et al., 2021).

Siderophore plates followed the same procedure as swimming plates, using 1X CC medium. In order to ensure that all inoculations were performed at the same distance from each other, they were all done by dipping five pipette tips in a cell suspension with a multichannel pipette and inserting the pipette tips until the center of the agar, as described for swimming inoculations.

### Analyses on publicly available data

For the analysis of *in vitro* genome-wide fitness data of root colonization in WCS417, we used supplementary data from (Cole et al., 2017) and selected the genes in the *iol* locus in WCS417 (*iolR*, PS417_11845; *iolC*, PS417_11850; *iolD,* PS417_11855; *iolE,* PS417_11860; *iolB*, PS417_11865; *iolI*, PS417_11870; *iolD*, PS417_11875; *iolG,* PS417_11880; *mocA*, PS417_11885; *iatA,* PS417_11890; *iatC*, PS417_11895; *iatB* PS417_11900). For the analysis of genome-wide fitness data of WCS417 in different conditions, we retrieved the data from (Price et al., 2018) available in Fitness Browser (https://fit.genomics.lbl.gov/cgi-bin/help.cgi). We kept all experimental conditions in which any of the genes in the *iol* locus of WCS417 was affected with a *t*-like score and a fitness value greater than 0.5. For the remaining conditions with more than one repeat, we took the median fitness value. We built a bipartite network of genes and conditions according to the fitness values using the *network* (Butts, 2008) and *ggnet* (Briatte, 2023), R packages for construction and visualization, respectively. For the network representation, we used the Fruchterman-Reingold algorithm with 1,000 iterations and an area and repulsion radius of 30,000,000. For the analysis of transcriptional activity of CHA0 in the host, we retrieved sequenced data from (Vesga et al., 2020) from the NCBI database (BioProject accession PRJNA595077) and mapped the reads to the CHA0 genome (assembly GCF_900560965.1) with kallisto v0.45.0 (Bray et al., 2016). Transcripts-per-million values for each gene were log_2_-transformed and scaled per gene.

## Supplementary Figures

**Figure S1. Schematic representation of the experimental design**. Root-associated pseudomonads are mixed through natural soil and let to colonise Arabidopsis roots for 21 days. Rhizosphere and root samples are collected and mixed with DNA of *Salinibacter ruber* as internal control. Absolute population density of each pseudomonad is calculated by sequencing the *16S rRNA* gene and normalising the counts with the internal control and the sample weight.

**Figure S2. The isolates in this study are a diverse set of root-associated pseudomonads**. (a) Venn diagram showing the relationship of ortholog group presence-absence among isolates’ genomes. Out of 8,752 orthogroups, 3,090 (35%) are shared by all isolates. (b) Distribution of the non-core genes. (c) Pair-wise average nucleotide identity (ANI) among isolates. (d) Pair-wise unweighted Jaccard similarity of the orthogroup presence-absence per genome (i.e., percentage of shared genes between isolate pairs). (e) Pair-wise unweighted Jaccard similarity of KEGG pathway annotations per genome. The relatively high values suggest that the functional capabilities of the isolates are very similar. (f) Same as e, but showing the weighted Jaccard similarity between the compositions of pathway annotations. The differences in the weighted vs unweighted metric indicate that certain functions encoded in the genomes do not distribute evenly between them, highlighting the functional diversity in the genomes. (g) Pair-wise correlation plots of the previously shown metrics in c-f.

**Figure S3. Microbial densities in inoculated and uninoculated samples.** Absolute inoculum densities at 21 dpi measured as the number of *16S rRNA* copies per gram of sample found in bulk soil (left), rhizosphere (middle), and root samples (right), for both inoculated and uninoculated samples.

**Figure S4. The isolates differ in their most colonized fraction.** Absolute inoculum densities after 21 dpi measured as the number of *16S rRNA* copies per gram of sample found in bulk soil (left), rhizosphere (middle), and root samples (right). *P. protegens* CHA0 and *P. simiae* WCS417 are highlighted in cyan and yellow, respectively, to differentiate them from the remaining isolates. The density of some isolates decreases in the root (RS158, WCS317, and WCS358), while others increase (WCS417, CHA0, and WCS134).

**Figure S5. Inositol effects on WT cells are highly variable.** Growth curves of WT and Δ*iol* cells in CC medium unsupplemented or supplemented with 10 mM glucose or inositol. The figure shows two examples of independent experiments. The effects of inositol in the WT are clearly variable per experiment, and the benefit of inositol on growth is only apparent in some cases.

**Figure S6. Inositol represses swarming motility in mutant cells.** (a) Swarming surface of WT (left) and Δ*iol* cells (right) in CC medium either unsupplemented or supplemented with glucose or inositol at 1 or 10 mM. Inositol does not seem to affect swarming motility in WT cells, while swarming is abolished in the mutant. (b) Same data as in a shown per treatment and pairing the swarming surface for both genotypes per plate, showing the difference in motility.

**Figure S7. Inositol affects mutant physiology.** Qualitative assessment of the effect of inositol and glucose on WT and mutant cells. Images show the same plates as in Figure 6 at 1, 2, and 3 dpi. Per day, the figure shows the plates organized as follows. First column: control conditions; second column: glucose 10 mM; third column: inositol 10 mM; first row: WT cells only; second row: combination of WT and mutant cells (WT, Δ*iol*, WT, Δ*iol*, WT); third row: mutant cells only. Mutant colonies start fading after reaching the end of the plate at approximately the end of the second day.

## Supplementary Tables

**Table S1:**
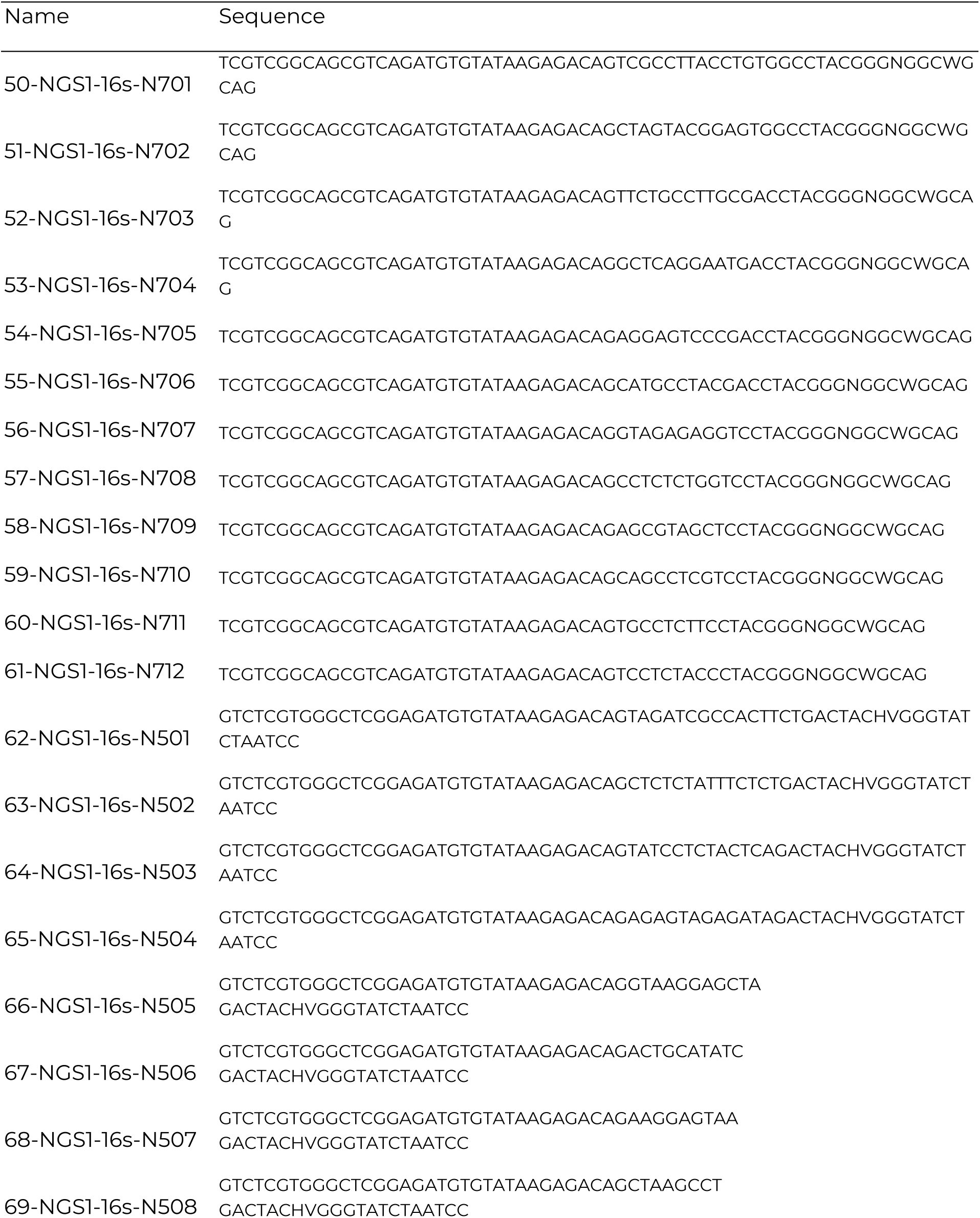
List of phasing primers used in this study for amplicon sequencing.

**Table S2:**
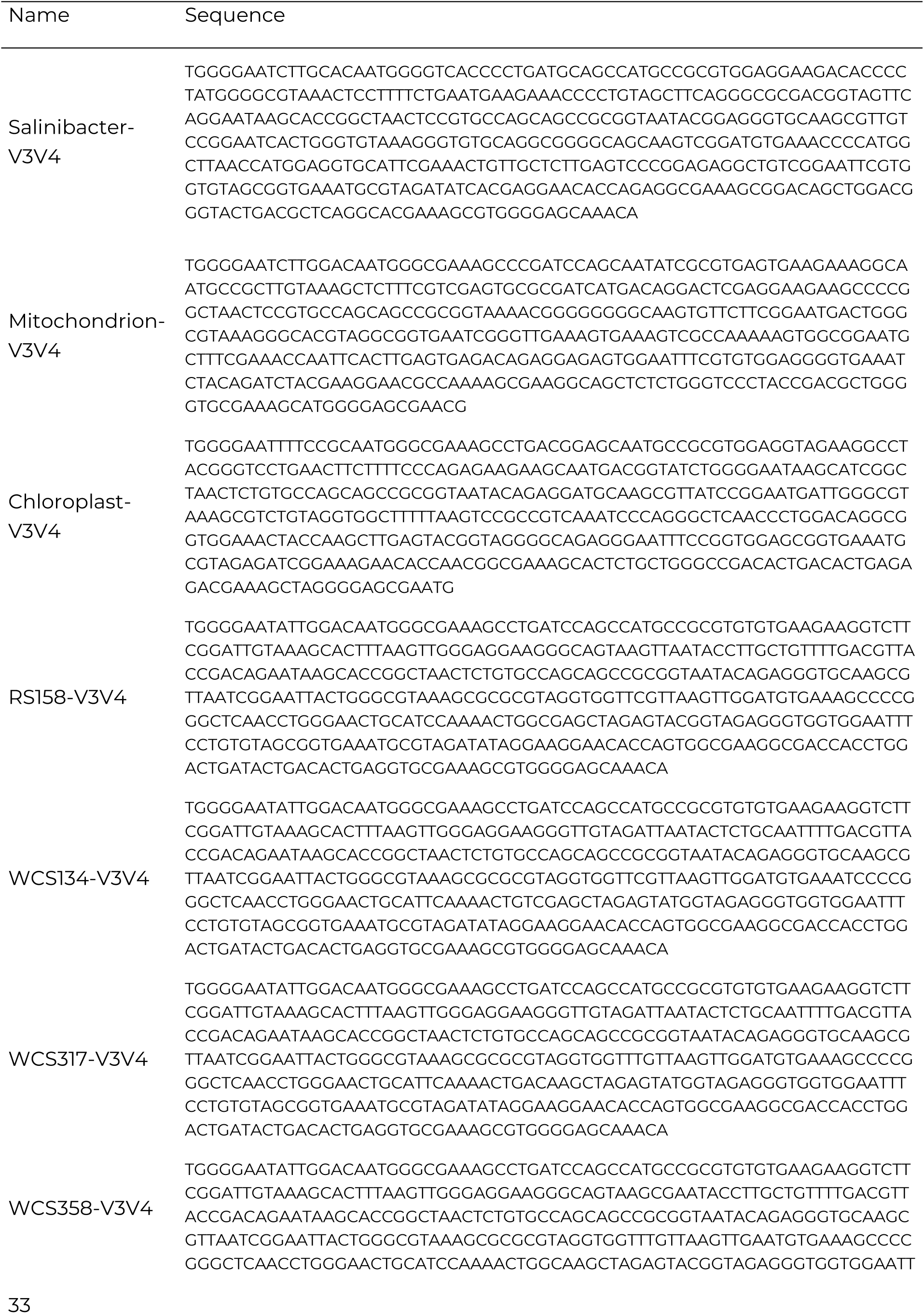
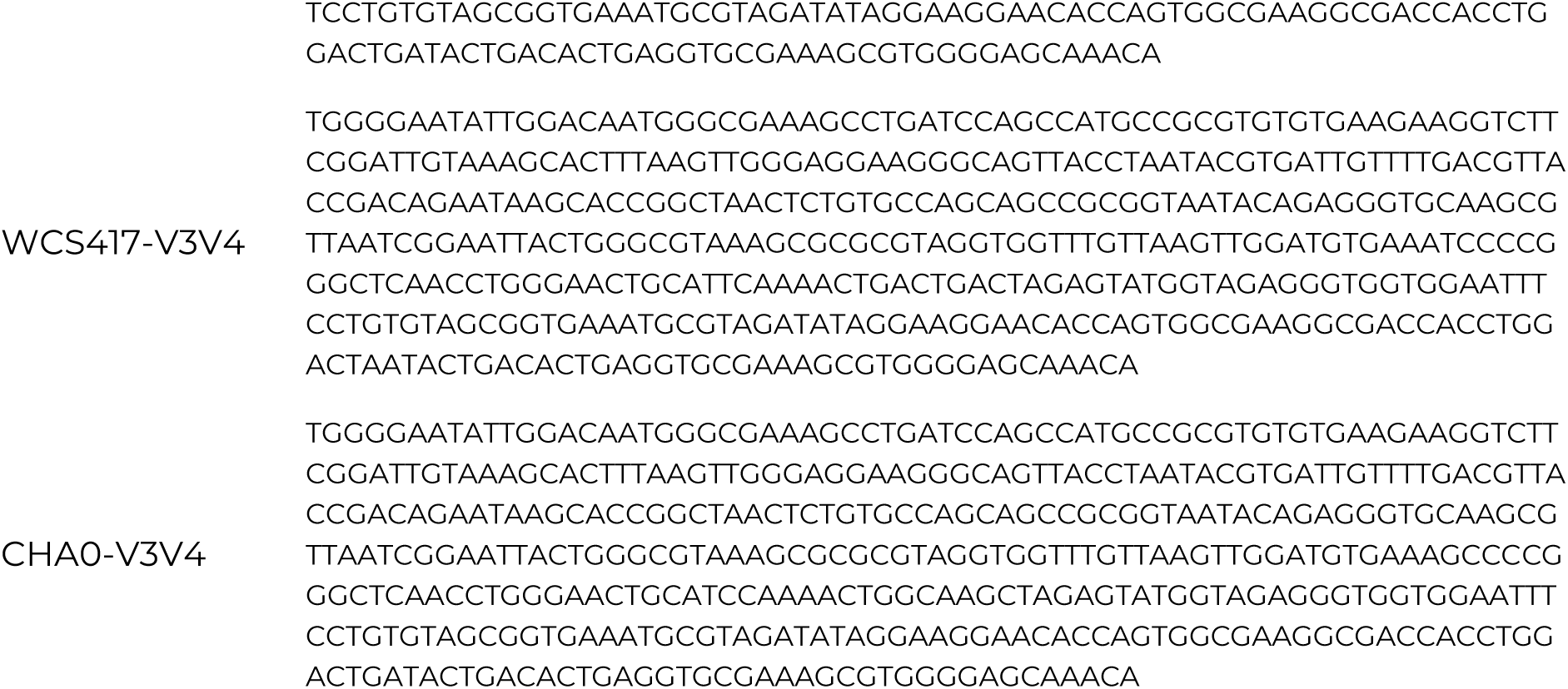
V3-V4 regions used as priors in the DADA2 pipeline.

**Table S3:**
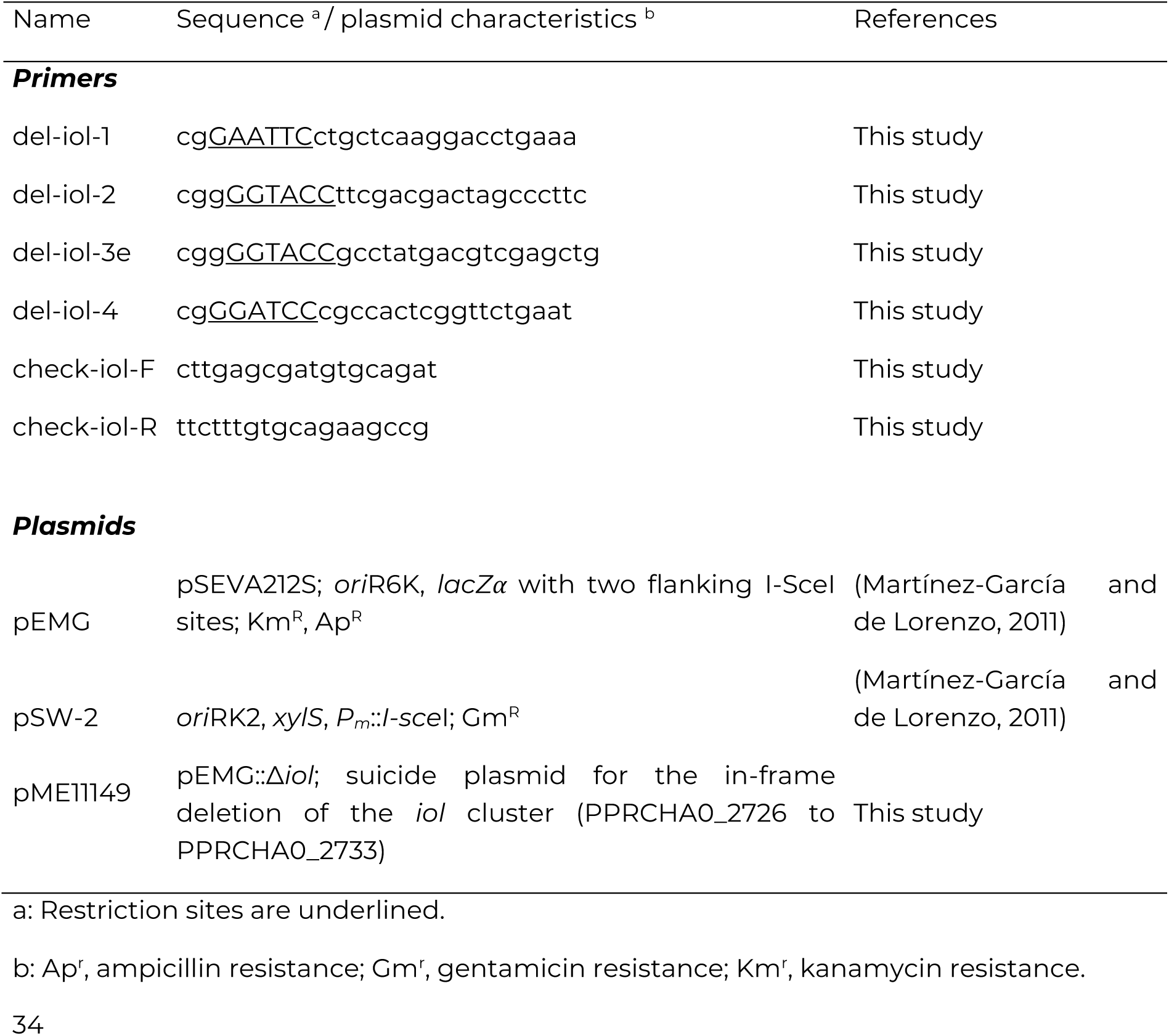
List of primers and plasmids used in this study for CHA0 mutagenesis.

