## Supplemental Information for "Identification of the conserved *iol* gene cluster involved in rhizosphere competence in *Pseudomonas*"

**Figure S7. Inositol affects mutant physiology.** Qualitative assessment of the effect of inositol and glucose on WT and mutant cells. Images show the same plates as in Figure 6 at 1, 2, and 3 dpi. Per day, the figure shows the plates organized as follows. First column: control conditions; second column: glucose 10 mM; third column: inositol 10 mM; first row: WT cells only; second row: combination of WT and mutant cells (WT, Δ*iol*, WT, Δ*iol*, WT); third row: mutant cells only. Mutant colonies start fading after reaching the end of the plate at approximately the end of the second day.

**Figure S1**

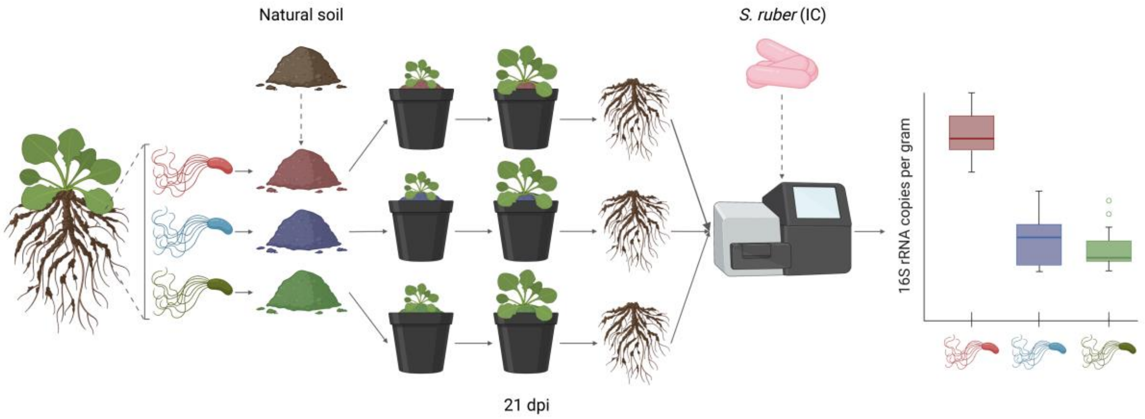

**Figure S2**

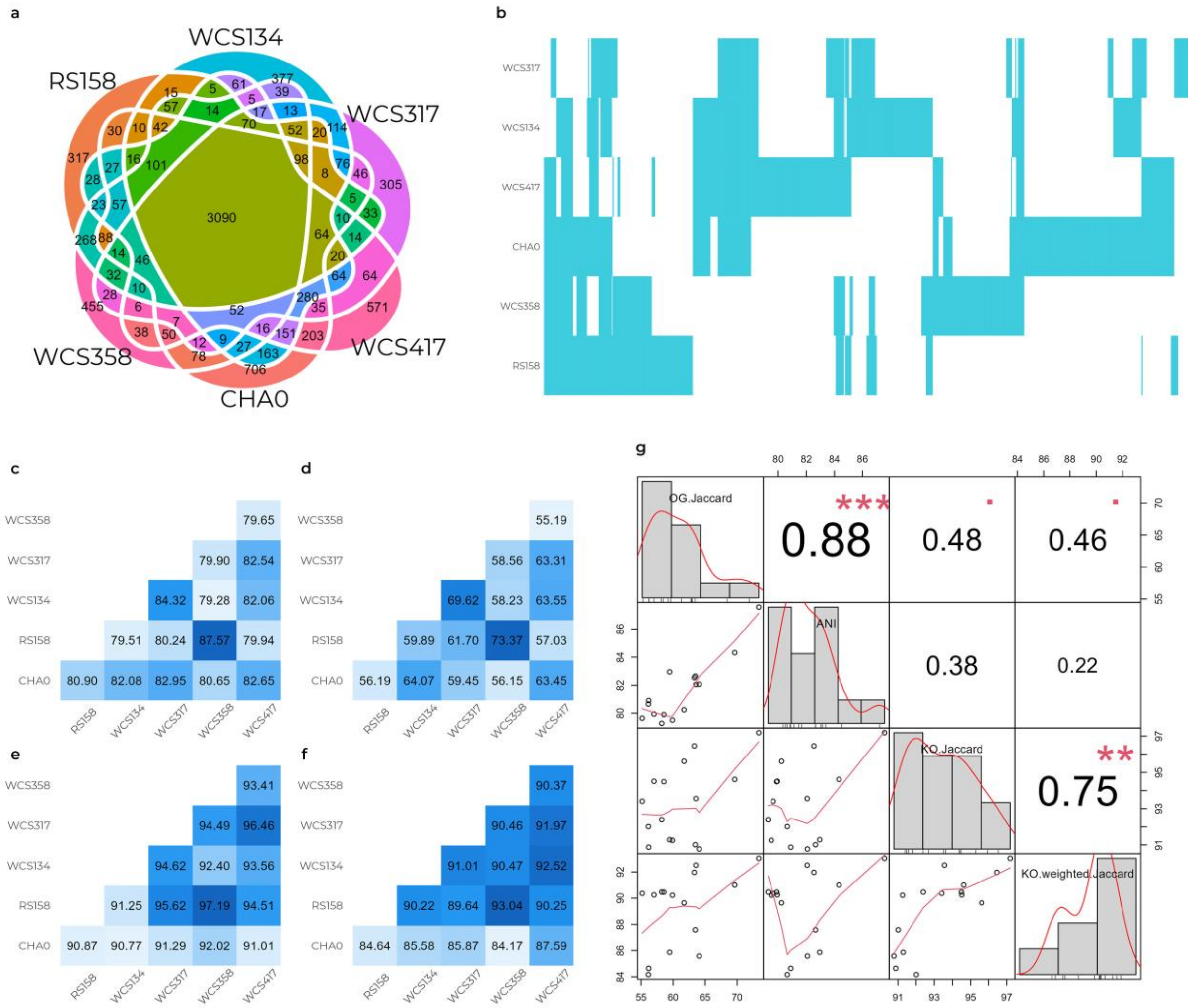
a
WCS134
RS158
WCS317
WCS417
WCS358
CHA0
b
WCS317
WCS134
WCS417
CHA0
WCS358
RS158
c
WCS358 79.65
WCS317 79.90 82.54
WCS134 84.32 79.28 82.06
RS158 79.51 80.24 87.57 79.94
CHA0 80.90 82.08 82.95 80.65 82.65
RS158 WCS134 WCS317 WCS358 WCS417
d
WCS358 55.19
WCS317 58.56 63.31
WCS134 69.62 58.23 63.55
RS158 59.89 61.70 73.37 57.03
CHA0 56.19 64.07 59.45 56.15 63.45
RS158 WCS134 WCS317 WCS358 WCS417
e
WCS358 93.41
WCS317 94.49 96.46
WCS134 94.62 92.40 93.56
RS158 91.25 95.62 97.19 94.51
CHA0 90.87 90.77 91.29 92.02 91.01
RS158 WCS134 WCS317 WCS358 WCS417
f
WCS358 90.37
WCS317 90.46 91.97
WCS134 91.01 90.47 92.52
RS158 90.22 89.64 93.04 90.25
CHA0 84.64 85.58 85.87 84.17 87.59
RS158 WCS134 WCS317 WCS358 WCS417
g
OG.Jaccard
ANI
KO.Jaccard
KO.weighted.Jaccard
*** 0.88
0.48
0.46
0.38
0.22
** 0.75

**Figure S3**

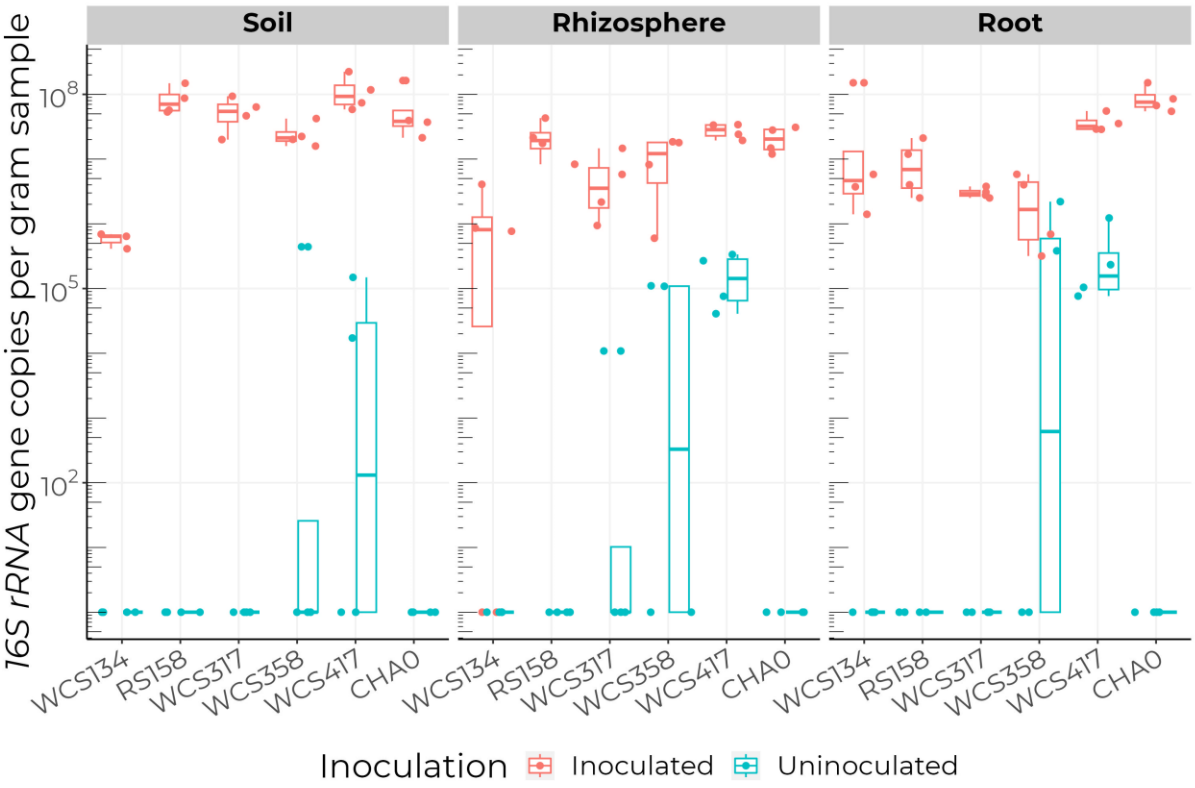
Soil
Rhizosphere
Root
16S rRNA gene copies per gram sample
10^8
10^5
10^2
WCS134
RS158
WCS317
WCS358
WCS417
CHA0
Inoculation
Inoculated
Uninoculated

**Figure S4**

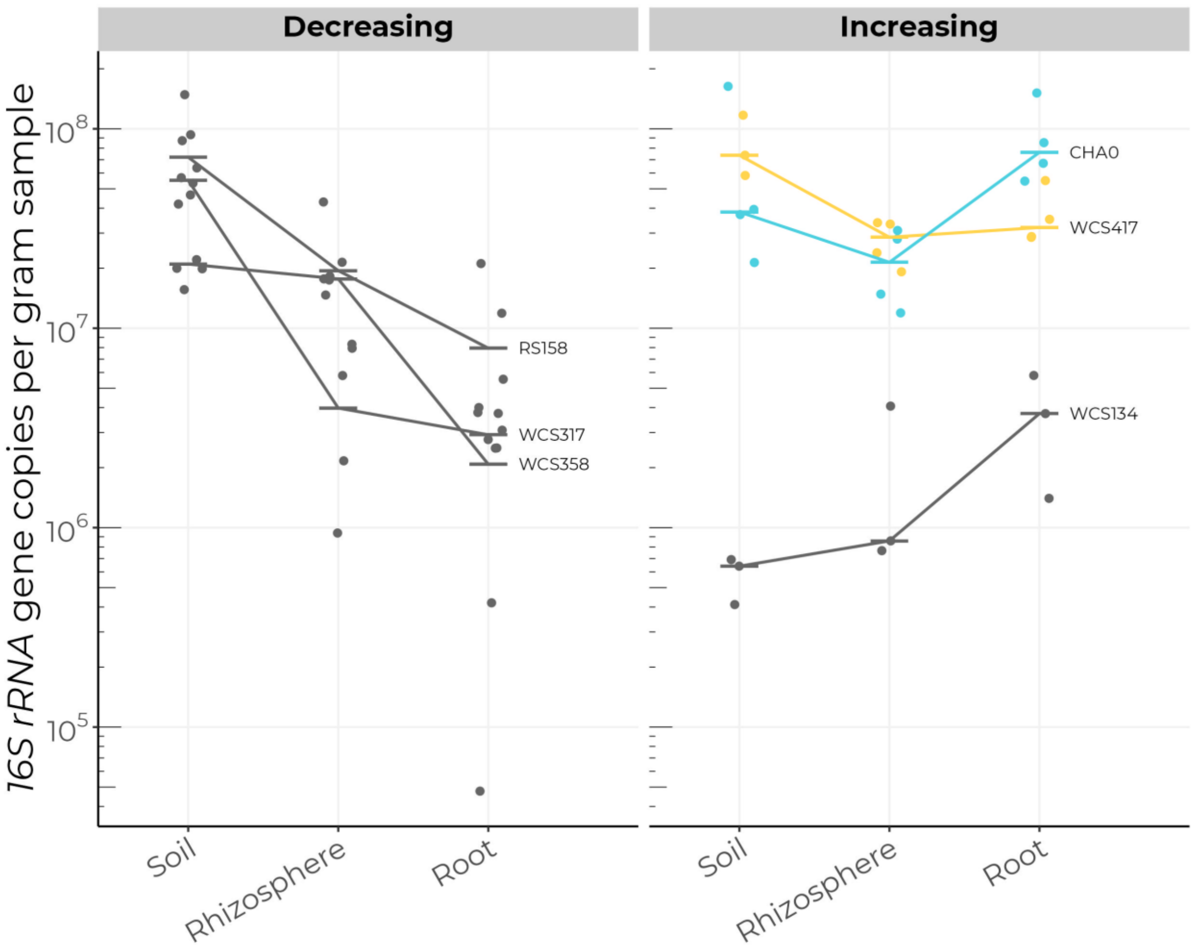
Decreasing
Increasing
16S rRNA gene copies per gram sample
10^8
10^7
10^6
10^5
RS158
WCS317
WCS358
CHA0
WCS417
WCS134
Soil
Rhizosphere
Root
Soil
Rhizosphere
Root

**Figure S5**

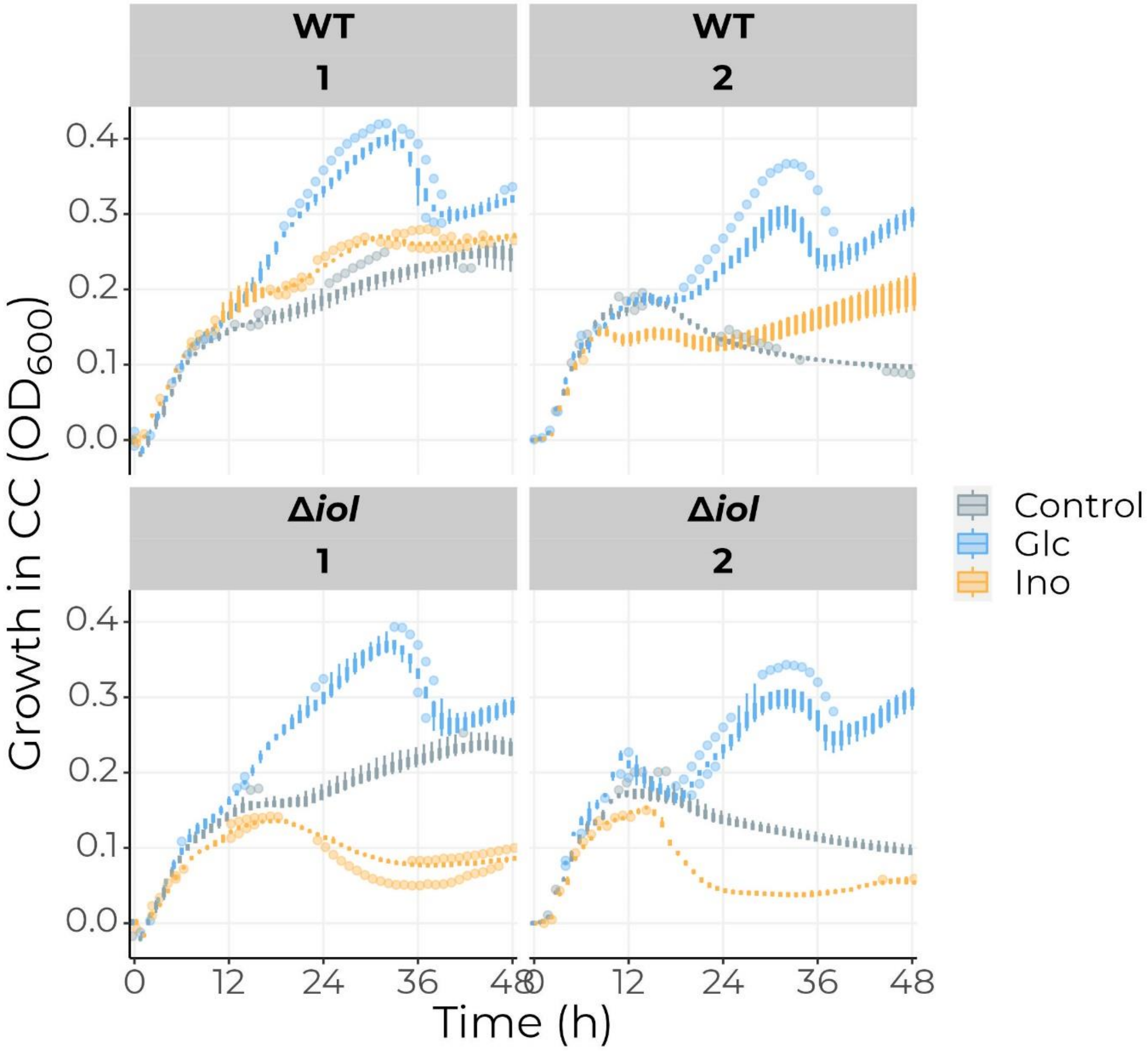
WT
1
WT
2
Δ*iol*
1
Δ*iol*
2
Growth in CC (OD600)
0.4
0.3
0.2
0.1
0.0
0
12
24
36
48
Time (h)
Control
Glc
Ino

**Figure S6**

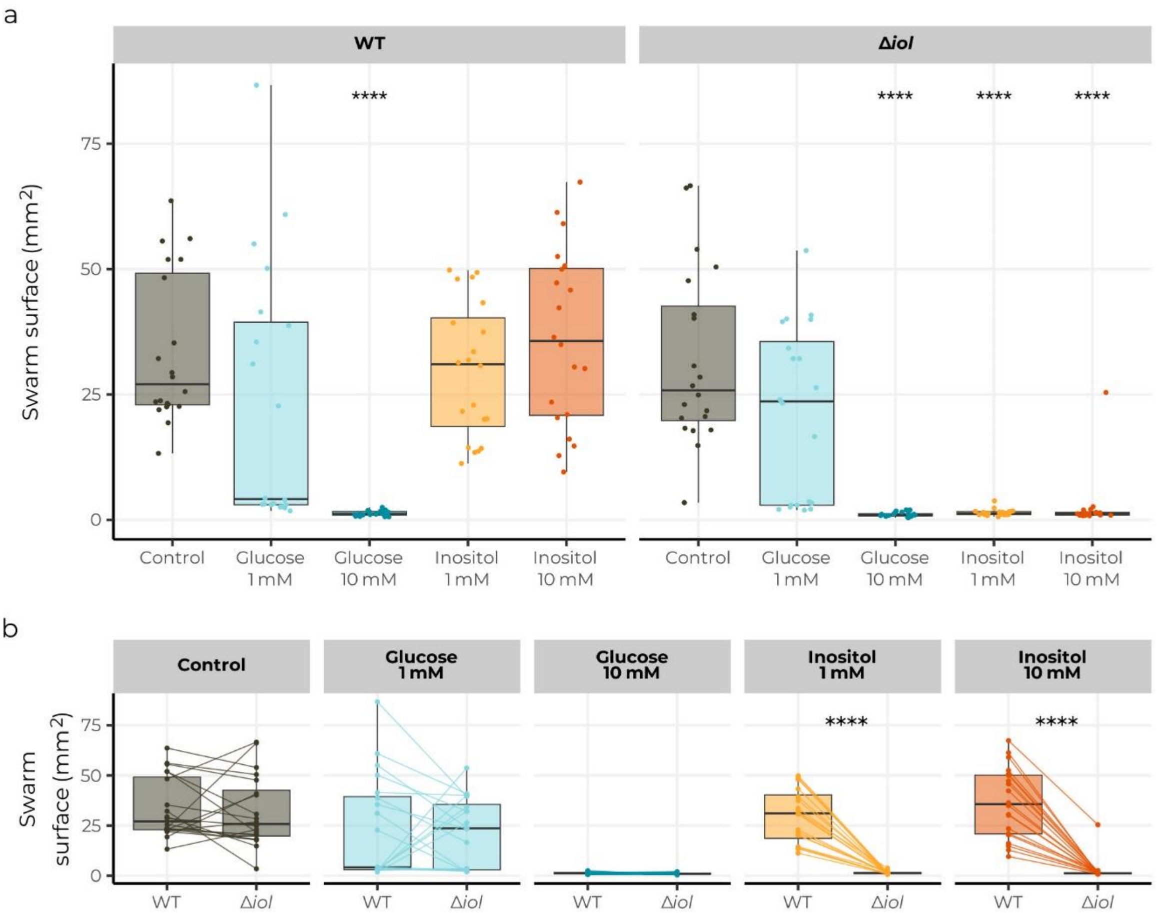
a
WT
Δiol
Swarm surface (mm²)
0
25
50
75
****
****
****
****
Control
Glucose 1 mM
Glucose 10 mM
Inositol 1 mM
Inositol 10 mM
b
Control
Glucose 1 mM
Glucose 10 mM
Inositol 1 mM
Inositol 10 mM
Swarm surface (mm²)
****
****
WT
Δiol

**Figure S7**

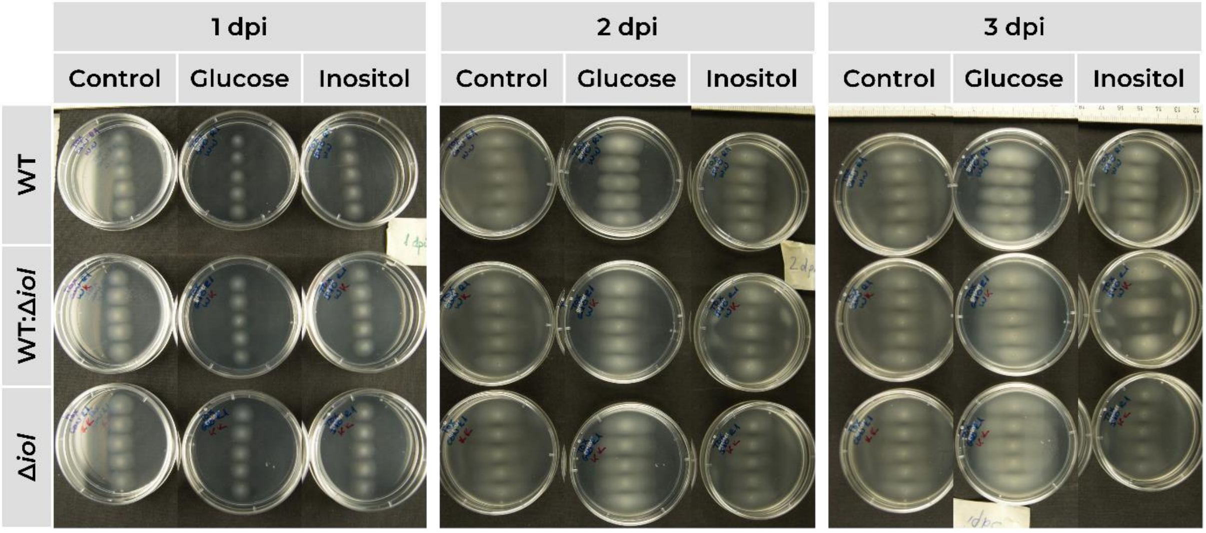
1 dpi
2 dpi
3 dpi
Control
Glucose
Inositol
Control
Glucose
Inositol
Control
Glucose
Inositol
WT
WT:Δ*iol*
Δ*iol*
